# Mice with impaired synaptic facilitation exhibit deficits in cognitive flexibility

**DOI:** 10.64898/2026.08.06.743101

**Authors:** Chloé L. Le Moing, Anna M. Bowman, Milana Krush, Joseph Gordon, Aaliya Mehnaz Ahmed, Skyler L. Jackman

## Abstract

Behavioral flexibility is crucial to animal survival in dynamic environments, and a failure to update actions in response to recent outcomes is a hallmark of many neuropsychiatric disorders. However, the cellular and circuit mechanisms in the brain that support behavioral flexibility remain poorly understood. Forms of short-term plasticity such as synaptic facilitation have been theorized to transiently maintain information in neural circuits, and flexibly modulate how circuits process information depending on recent activity. Despite theoretical support, there is no direct experimental evidence linking synaptic facilitation to flexible decision-making. Recently, the presynaptic calcium sensor Synaptotagmin-7 (*Syt7*) has been shown to be required for synaptic facilitation at many synapses in the mammalian brain. Here, we assess operant learning in male and female *Syt7* KO mice to determine how facilitation contributes to learning both stable and dynamic reward contingencies. We find that *Syt7* KO mice acquired stable contingencies similarly to wild-type controls. However, KO mice were impaired in learning dynamic contingencies, showed more perseverative responding, and were delayed in applying a new task rule to all trial types following reversal. Behavioral modeling revealed a reduced influence of recent trial history on decisions in KO mice compared to wild-type controls. The behavioral deficits could not be explained by differences in motivation or memory. These results suggest that synaptic facilitation supports adaptive decision-making and that disruptions of short-term plasticity impair animals’ ability to use recent outcomes to update behavior.

## Introduction

In a world defined by constant change, survival depends not only on learning, but on the ability to unlearn. Food sources disappear, threats emerge unexpectedly, and actions that once led to reward may suddenly produce failure. To navigate these shifting contingencies, animals must flexibly update stimulus-response associations, revise behavioral strategies, and continually reevaluate outcomes. Impairments in these processes are core features of many neuropsychiatric disorders, including autism spectrum disorder, schizophrenia, and Alzheimer’s disease (Uddin, 2021).

A well-studied form of flexible behavior is reversal learning, in which a subject learns a stimulus-or action-outcome contingency and must subsequently adapt to respond correctly when the contingency has reversed. At the neural level, flexibility depends on the ability of cortical and subcortical circuits to update neural representations of task-relevant information based on recent experience and adjust behavior accordingly (Miller and Cohen, 2001; Tervo et al., 2014; Stokes et al., 2017). These computations require neural activity to be shaped not only by current inputs, but also by information carried forward across short timescales. A growing body of theoretical work has therefore proposed that short-term synaptic plasticity (STP) could support flexible behavior by transiently storing recent information and modulating how inputs are integrated over time (Mongillo et al., 2008; Stokes et al., 2013; Aitken and Mihalas, 2023).

Despite strong theoretical support from decades of computational modeling studies, there remains no direct evidence linking STP to flexible behavior. Techniques for observing STP in behaving animals, including directly via intracellular recordings and indirectly via extracellular recording, are difficult and generally low-yield (Fujisawa et al., 2008; Tao et al., 2015). However, the identification of protein machinery responsible for STP in recent decades allows for genetic manipulations and characterization of subsequent behavioral phenotypes.

Synaptotagmin-7 (SYT7) is one such molecular target. It is a presynaptic calcium sensor required for synaptic facilitation in a wide range of brain regions, including cortex, hippocampus, thalamus, and cerebellum (Jackman et al., 2016; Chen et al., 2017; Martinetti et al., 2022; Chiu and Carter, 2024; Shin et al., 2026). Though synaptic phenotypes in *Syt7*-knockout (KO) neurons have been extensively characterized, behavioral and neurological phenotypes linked to SYT7 are far less studied. Genome-wide association studies have identified genetic loci linked to cognitive performance, educational attainment, and intelligence metrics that implicate *Syt7* as a candidate gene (Lee et al., 2018; Cerezo et al., 2025).

SYT7 has further been implicated in a variety of conditions with cognitive symptoms, including Alzheimer’s disease, bipolar disorder, and multiple sclerosis (Barthet et al., 2018; Fritsche et al., 2020; Shen et al., 2020). Independent tissue enrichment and transcriptomic analyses across humans and animals show that *Syt7* expression is enriched in cerebral cortex, consistent with a potential role in cortical circuits supporting higher cognitive functions (Lein et al., 2007; Shen et al., 2012; Wu et al., 2023).

Recent studies have reported a role for SYT7 in affective behavior, fear conditioning, and pattern completion (Shen et al., 2020; Xie et al., 2021; Marneffe et al., 2024), but the role of SYT7-driven facilitation in operant learning and flexibility remains unknown. We analyzed performance and decision-making in *Syt7* KO mice to probe the effect of facilitation deficits on acquisition and reversal learning.

KO animals showed no deficits in acquisition of short-term memory tasks with stable contingencies, but had impaired performance when reward contingencies were not stable. Behavioral modeling on a probabilistic task with multiple reversals revealed an impairment in the ability to use recent outcomes to update ongoing behavior. Our results suggest that synaptic facilitation plays a key role in reversal learning but is dispensable for learning action-outcome associations.

## Materials & Methods

### ANIMALS

All animal procedures were performed in accordance with protocols approved by the Institutional Animal Care and Use Committee of [Author University]. Adult (2-6 months) male and female *Syt7* KO mice (B6.129S1-*Syt7*tm1Nan/J, Jax:004950, https://www.jax.org/strain/004950) and wild-type littermate controls were used in all experiments. Mice were housed in groups of two to five on a 12-hr reversed light-dark cycle (ZT 12 / lights off at 12pm), and experiments were conducted under red or dim white lighting during the first 1-5 hours of the dark cycle. All operant assays and the Y-maze assay took place in an area of the animal colony housing room designated for experiments, eliminating the need for animals to be transported or habituated prior to experimentation. Open field and rotarod assays were performed in a separate room, to which animals were habituated >1 hr before testing. All animals were habituated to handling for three consecutive days prior to experimentation. Behavioral equipment was cleaned with unscented dish soap or ethanol between animals to reduce odor contamination. Animals used in operant assays were only trained on a single task, except where noted. All assays were performed by experimenters blinded to animal genotypes. Mice were weighed prior to experimentation to obtain baseline weights then food-restricted (radial arm maze) or water-restricted (all other operant assays) to 85-90% of their baseline weight so that food or water rewards served as effective reinforcers during operant procedures. Weights were monitored daily immediately before experimental sessions for the duration of testing, and target weights were increased by 0.5g weekly to account for normal growth.

Food-restricted mice received water ad libitum and 50-70% of their daily ad lib food consumption (estimated as 15g of food per 100g body weight), delivered at least 1 hour following experimental sessions. Water-restricted mice received food ad libitum and were able to achieve up to 800μL of water daily during training, with 1-2% citric acid water ad libitum on weekends to maintain restriction (Urai et al., 2021). Any animal falling below 85% of its baseline weight or failing to obtain the target amount of water during operant training received additional food or water to meet daily requirements.

### OPERANT TASKS

Operant training was conducted using Bpod operant chambers with protocols controlled by Bpod State Machines (Sanworks LLC, Stony Brook, NY). In all operant procedures, the entire apparatus was contained in a custom sound-isolating box. Each chamber contained either five adjacent water-dispensing nose-poke ports on one wall and one on the opposite wall (5-port delayed match-to-sample task) or three adjacent ports on one wall (all other operant tasks). Ports could be illuminated and dispense measured water rewards when nose-poked, as detected via infrared beam breaks. Illumination of a bright overhead house light served as a punishment cue for unsuccessful trials. The tasks were programmed with custom MATLAB code. All responses during tasks were recorded and saved in MATLAB.

Similar initial training protocols were employed for all behavioral tasks. During training, correct pokes triggered immediate 10-µL reward delivery and a 1-to 3-second inter-trial interval (ITI) with all lights extinguished. Incorrect choice pokes triggered illumination of the house light and a five-to-ten second ITI, serving as behavioral feedback for unrewarded or “punish” trials. Failure to initiate a trial or make a choice within the limited hold window of 10-20 seconds was registered as an omission and triggered a trial restart. The criterion for successful learning was for mice to perform at 75% accuracy for three consecutive sessions, with accuracy calculated as number of correct trials ÷ number of incorrect trials. All sessions were limited to one hour or 500 trials. Training was performed in stages to shape behavior toward the end task.

Spatial short-term memory and reversal learning were assessed using a match and non-match to sample protocols adapted from Yhnell et al. (2016). For match/non-match reversal learning, mice were first trained for 12 sessions to perform a match-to-sample task, followed by an unsignaled switch to a non-match to sample for an additional 8 sessions. Each trial began when the mouse initiated the trial by nose-poking into the center port, triggering the illumination of either the left or right port (sample phase). A nose-poke into the illuminated sample port was required, followed by a nose-poke into the center, after which both the left and right ports would illuminate (choice phase). The mouse was required to select the same port previously visited during the sample phase to receive a reward for the match task, or the port not previously visited for the non-match task. The initiation poke requirement was removed from the non-match reversal task, and trials began with the sample phase. The primary measures of interest to assess reversal learning were proportion of correct responses, perseverative and regressive error rates, choice and reward poke latencies, proportions of win-stay and lose-stay choices, and side bias indices during the non-match stage. The total correct proportion of each trial type across all consecutive sessions was calculated for each mouse, and the trial types were classified as a high-performance and low-performance trial type for comparing error types. Error types were calculated for each trial type (right-sample and left-sample) separately. Perseverative errors were calculated as incorrect choices made before a reaching a criterion of 35 cumulative correct choices, and regressive errors were calculated as incorrect choices made after this criterion. Side bias index was calculated as the absolute difference in accuracy between left-and right-sample trials.

In the 3- and 5-port match-to-sample tasks with enforced delays (Fig. 3), the training protocol was identical except that choice lights would not illuminate until a predetermined delay had elapsed (0, 2, 5, 10, or 20 seconds) after the delay poke. Mice were trained on the 3-port task for 10-12 sessions, and a subset was further trained on the 5-port version of the same task for 9-10 sessions of shaping and 11 sessions of the full task. The primary measure of interest to assess short-term memory was the proportion of correct responses for each delay length and sample location.

Cognitive flexibility was assessed using a probabilistic serial reversal learning task adapted from Choi et al. (2023). Each trial began when the mouse nose-poked into a center port to initiate the trial. Both side ports were then illuminated, and the mouse was required to choose one port. One side was designated as the high-probability reward port (85% chance of reward), while the other was a high-probability punishment port (85% chance of house light). Both ports had a 15% chance of no outcome. After eight reward port choices out of ten consecutive trials, the contingencies were reversed. Reversals continued throughout the session, requiring mice to adapt to the changing rules. The initial correct port (left or right) was alternated between each of the 8 sessions. The primary measures of interest to assess cognitive flexibility were the proportion of reward port choices and the number of trials per reversal.

Behavioral modeling was performed using code from Choi et al. (2023) available on GitHub (github.com/Fuccillo-Lab/Choi_et_al_2023_NatComm). A win-stay/lose-shift model was used to estimate the tendency of mice to lapse from the optimal strategy of repeating rewarded choices and switching after punished choices, with parameter ε representing the lapse rate as the probability of win-shift/lose-stay choices. A logistic regression model was used to predict the probability of choosing the righthand port 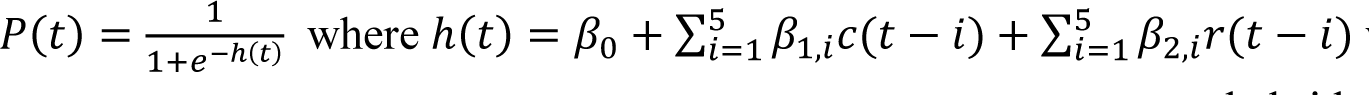 where *c(t)* is the choice on trial *t* coded as +1 for right and -1 for left, and *r(t)* represents the rewarded side as +1 if a reward came from a right choice, -1 for a left choice, and 0 for no reward (Choi et al., 2023).

### RADIAL ARM MAZE

The maze used in this task comprised eight 34cm-long 5cm-wide arms made of opaque plastic projecting from a 20-cm diameter center and was placed on a table in a room with several fixed visual cues. During testing, each arm was baited with one highly palatable, odor-free reward pellet (Bio-Serv Dustless Precision Pellets) in a win-shift task. Plastic food cups placed at the end of each arm allowed for the arms to be baited without rewards being visible to the animal while traversing the maze. Prior to testing, mice were habituated to handling and reward pellets. Mice were gradually habituated to the maze over three days. On the first day, cage mates were placed in the maze together with scattered reward pellets for 15 minutes under red light. This was repeated the following day without cage mates for five minutes in red light followed by five minutes in dim white light. On the final day of habituation, each mouse was placed in the fully-baited maze for 7.5 minutes under dim white light. During testing, each mouse was confined in the center of the maze for 10 seconds before being released to freely explore the arms. A session ended when a mouse had entered all eight arms once or when five minutes elapsed. An arm was recorded as visited if the animal traveled at least half the distance to the food cup (17 cm). Testing was repeated daily for 11 days with one rest day at the halfway point. After an additional rest day, this protocol was repeated for four more days with a slightly altered protocol, in which mice were confined to the center for ten seconds after each arm entry to prevent the use of mediation strategies (e.g. sequentially entering adjacent arms). The primary measures of interest to assess spatial short-term/working memory were the number of arms visited before repeat and total number of repeat entries (errors).

### ROTAROD

Mice were given three trials per day for three days. Trials started with a rotation speed of 0.5 rotations per minute (RPM) and increased incrementally by one RMP every three seconds. The time which a mouse fell was recorded for each trial and averaged each day.

### OPEN FIELD

Mice were placed in a round plastic 43-cm diameter circular arena and their behavior was recorded for five minutes. Behavior was recorded to quantify locomotor activity and time spent in the center third of the chamber.

### Y-MAZE

The maze used in this task comprised three 30cm-long 10cm-wide arms made of opaque plastic projecting from a triangular center and was placed on a table in a room with several fixed visual cues. Spontaneous behavior was recorded for eight minutes. Behavior was recorded to quantify locomotor activity and spontaneous alternation.

### STATISTICAL ANALYSIS

All scoring and data analysis was performed blind to genotype using custom scripts written in MATLAB or Python. Video recordings of mice in the open field, radial arm maze, and Y maze were analyzed using ezTrack (https://github.com/denisecailab/eztrack) (Pennington et al., 2019). Distribution normality was checked with the Shapiro-Wilk or Kolmogorov-Smirnov test. T-tests or nonparametric equivalent Wilcoxon or Mann-Whitney tests were used to check significance. Statistical significance was set at p < 0.05. To assess behavior across multiple contexts (e.g. session number), two-way mixed ANOVA with Greenhouse-Geisser corrections were used with factors of genotype and context followed by post-hoc t-tests for cases where the residuals were normally distributed. If residuals were not normal, a linear mixed effects model with context and genotype as fixed effects and subject as a random intercept was used with post-hoc Wilcoxon or Mann-Whitney tests for paired and unpaired comparisons, respectively. Greenhouse-Geiser and Bonferroni corrections were applied where appropriate.

## Results

### Syt7 KO mice show impaired rule reversal learning

To assess the role of SYT7 in cognitive flexibility, we trained wild-type (WT) and *Syt7* knockout (KO) mice on a two-choice match-to-sample task, followed by an unsignaled reversal to a non-match rule (Fig. 1A). This design allowed us to compare initial acquisition under stable contingencies against performance following a change in task structure.

**Figure 1:**
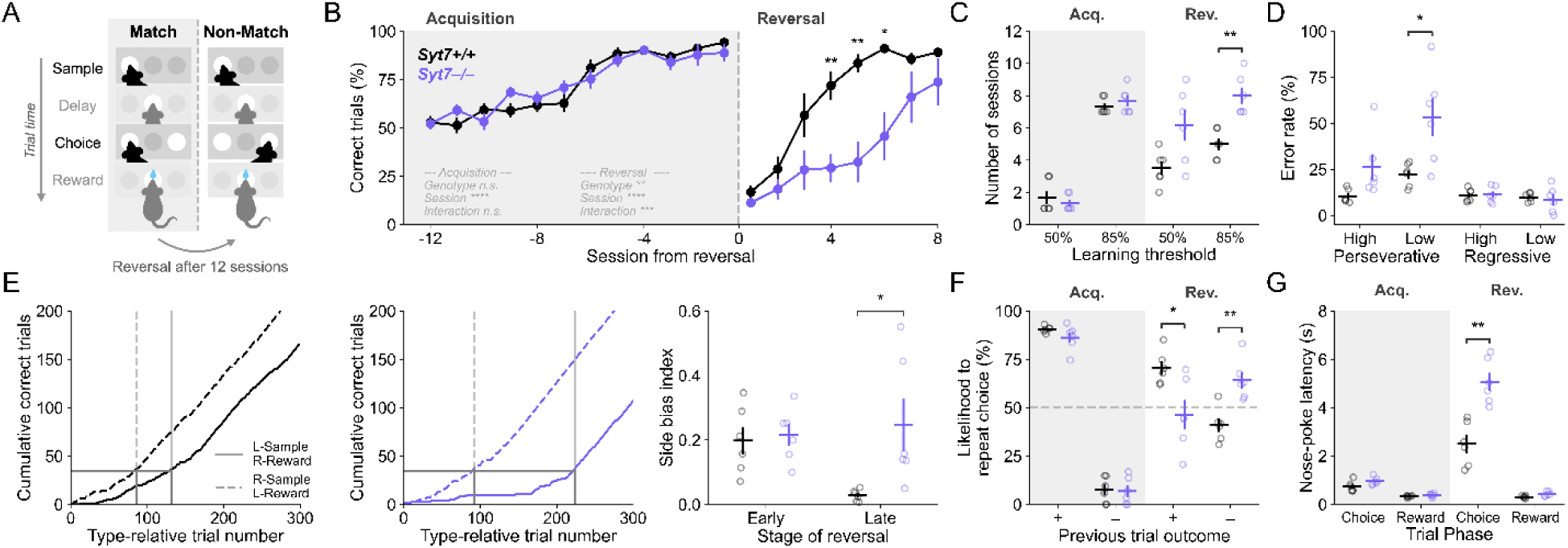
Knockout of Syt7 impairs match-nonmatch reversal learning but not acquisition learning. Shaded backgrounds indicate match-to-sample acquisition (“Acq.”) while white backgrounds indicate reversal to non-match (“Rev.”). A) Schematic of match to non-match reversal task. B) Proportion of correct trials across all training sessions for WT (N=6) and *Syt7* KO animals (N=6). Shaded area represents match-to-sample acquisition. Dotted line indicates rule change from match to non-match. C) Number of training sessions required to reach a session-level threshold of chance performance (50% of trials correct) and criterion performance (85% of trials correct). D) Perseverative and regressive errors during non-match. Error type proportions were calculated separately for right- and left-sample trials, then classified as higher- (“High”) or lower-performance (“Low”) trial types for each mouse. Perseverative errors were calculated as incorrect choices made before reaching a criterion of 35 cumulative correct, and regressive errors calculated as incorrect choices made after this criterion. E) Left: Exemplar learning curves for one WT (left) and one KO (right) mouse showing distinct learning curves for left(L)-sample right(R)-reward and R-sample L-reward trial types across all non-match trials in consecutive sessions. Vertical lines indicate learning criterion, calculated as the number of cumulative trials required to reach 35 total correct for each trial type. Right: Difference in trial type learning calculated as the absolute difference in the proportions of correct L-sample and correct R-sample trials, split by early (first 4) and late (5^th^ onward) non-match sessions. F) Repeat choices following rewarded (+) and punished (–) trials quantified individually for each trial type and averaged across sessions. G) Nose poke latencies for choice and reward pokes, as diagrammed in *A*. Data points are expressed as mean ± SEM, with small hollow circles representing individual animal replicates. Statistical significance is shown as *p<0.05, **p<0.01, ***p<0.001, and ****p<0.0001.

Both WT and KO mice acquired the initial match-to-sample task at similar rates, taking comparable numbers of sessions to reach chance (WT 1.7±0.42; KO 1.3±0.21) and criterion performance (WT 7.3±0.21; KO 7.7±0.33; Fig. 1B,C). In contrast, following reversal to the non-match rule, KO mice exhibited a marked impairment. Although both groups initially dropped to below-chance performance, WT mice rapidly adapted to the non-match rule, whereas KO mice required significantly more sessions to reach criterion (WT 5.0±0.37; KO 8.0±0.52; *t*(10)=-4.74; *p*=0.0031; Fig. 1B,C). This difference persisted when performance was analyzed as a function of trials to criterion rather than sessions (WT 286.5±30.3; KO 396.2±43.5; *U*=5, *p*=0.045), indicating that slower learning in KO mice was not explained by reduced task exposure (Extended Data Fig. 1-1A). Additionally, impaired performance of KO mice could not be explained by an inherent deficit in non-match learning, as a separate cohort of mice trained on non-match alone showed improvement across sessions (all *p*<1.7×10^-7^) with no significant differences between genotypes (genotype or interaction effect: all *F*<1.3, all *p*>0.3; Extended Data Fig. 1-1B).

Visual inspection of reversal-stage behavior suggested that mice did not perform equivalently across trial types. Instead, many animals displayed a pronounced preference for one trial type, leading to consistently higher accuracy on one trial type than the other. Because such biases may obscure the mechanisms underlying behavioral flexibility, we categorized trial types within each mouse as high-performance and low-performance and analyzed error patterns separately within these categories.

Perseverative errors reflect continued application of the previously rewarded match-to-sample rule and were more frequent in KOs than WTs, with this difference being statistically significant in the low-performance trial type (*t*(10)=-2.91, *p*=0.031; Fig. 1D). Regressive errors were instead low and consistent across genotypes, indicating a similarly infrequent return to the previously reinforced rule after having learned the new rule.

Consistent with this, KO mice exhibited more persistent asymmetries in performance across trial types than WTs, with higher side bias (calculated as the absolute difference in accuracy between left-and right-sample trials) in the late phase of reversal learning (*t*(10)=-2.65, *p*=0.0488; Fig. 1E). In a two-choice task, such asymmetries can arise either from incomplete application of the updated rule to both trial types or from a simpler strategy, such as a bias toward one choice port independent of which port was sampled (Zhu and Kuchibhotla, 2024). To examine how recent outcomes influenced choices, we analyzed win-shift/lose-stay strategies separately for each trial type. Although both groups showed similar strategies overall during match task acquisition, KO mice were less likely repeat rewarded choices (*t*(10)=2.90, *p*=0.032) and more likely to repeat unrewarded choices (*t*(10)=-4.24, *p*=0.0034) during the reversal to non-match (Fig. 1F).

The lower success rates of KO mice after rule reversal could reflect an impairment of flexible behavior, or it might be explained by non-specific factors such as impaired motivation or motor output. Although motivation and motor function have not been extensively characterized in *Syt7* KO mice, these animals exhibit mild hyperlocomotion during light/dark box tests of anxiety (Liu et al., 2025). To investigate these possibilities, we assessed the time it took animals to complete different phases of the task. Choice and reward latencies were analyzed as correlates of deliberation and motivation, respectively. Interestingly, although KO and WT mice exhibited similar choice latencies during the match rule, KOs showed significantly longer choice latencies than WTs after reversal to non-match (WT 2.5±0.4; KO 5.1±0.0.4; *U*=0, *p*=0.0087) with no genotype difference in reward poke latency in either stage (all *p*>0.1; Fig. 1G). This argues against major motor deficits, as both WT and KO mice were capable of reaching the reward port within 500ms on average regardless of learning stage. Further arguing against generalized motivation deficits, KO and WT animals completed similar numbers of trials while learning the match rule (Extended Data Fig. 1-1A).

Together, these results demonstrate that *Syt7* KO mice have a selective impairment in reversal learning. This deficit is not driven by reduced motivation or failure to maintain a learned rule, but instead reflects an impairment in using feedback to adjust actions appropriately following a change in task contingencies.

### Syt7 KO mice show reduced sensitivity to recent probabilistic outcomes

To determine whether SYT7 contributes to flexible behavior when outcomes are probabilistic, we trained WT and *Syt7* KO mice on a serial reversal task in which reward contingencies reversed dynamically based on recent performance and outcomes were not always delivered (Fig. 2A). This task requires animals to continuously update action values based on recent outcomes. KO mice exhibited impaired adaptation to changing contingencies: while overall task performance across sessions did not differ significantly between genotypes (Fig. 2B), KO mice required significantly more sessions to reach chance-level performance (WT 1.83±0.17; KO 3.33±0.49; *U*=5, *p*=0.025; Fig. 2C) and more trials to reach criterion following each reversal (WT 19.5±1.1; KO 26.0±1.7; *t*(10)=-2.23, *p*=0.0495; Fig. 2D). We again tested whether this impairment could be explained by reduced motivation or engagement. KO and WT mice initiated a similar number of trials per session (WT 236.4±15.7; *Syt7* KO 215.0±20.1; *t*(10)=0.84, *p*=0.42) and showed no differences in average initiation latency (WT 10.2 ±0.9; KO 12.6±10.2; *U*=12, *p*=0.34), suggesting comparable task engagement. However, the preceding trial outcome had a strong influence on the time it took WT mice to initiate the next trial, with shorter latencies following rewarded trials (*t*(5)=6.37, *p*=0.0042; Fig. 2E). This effect was attenuated in KO mice (*p*=1.0, corrected), suggesting a reduced behavioral sensitivity to reward outcomes.

**Figure 2:**
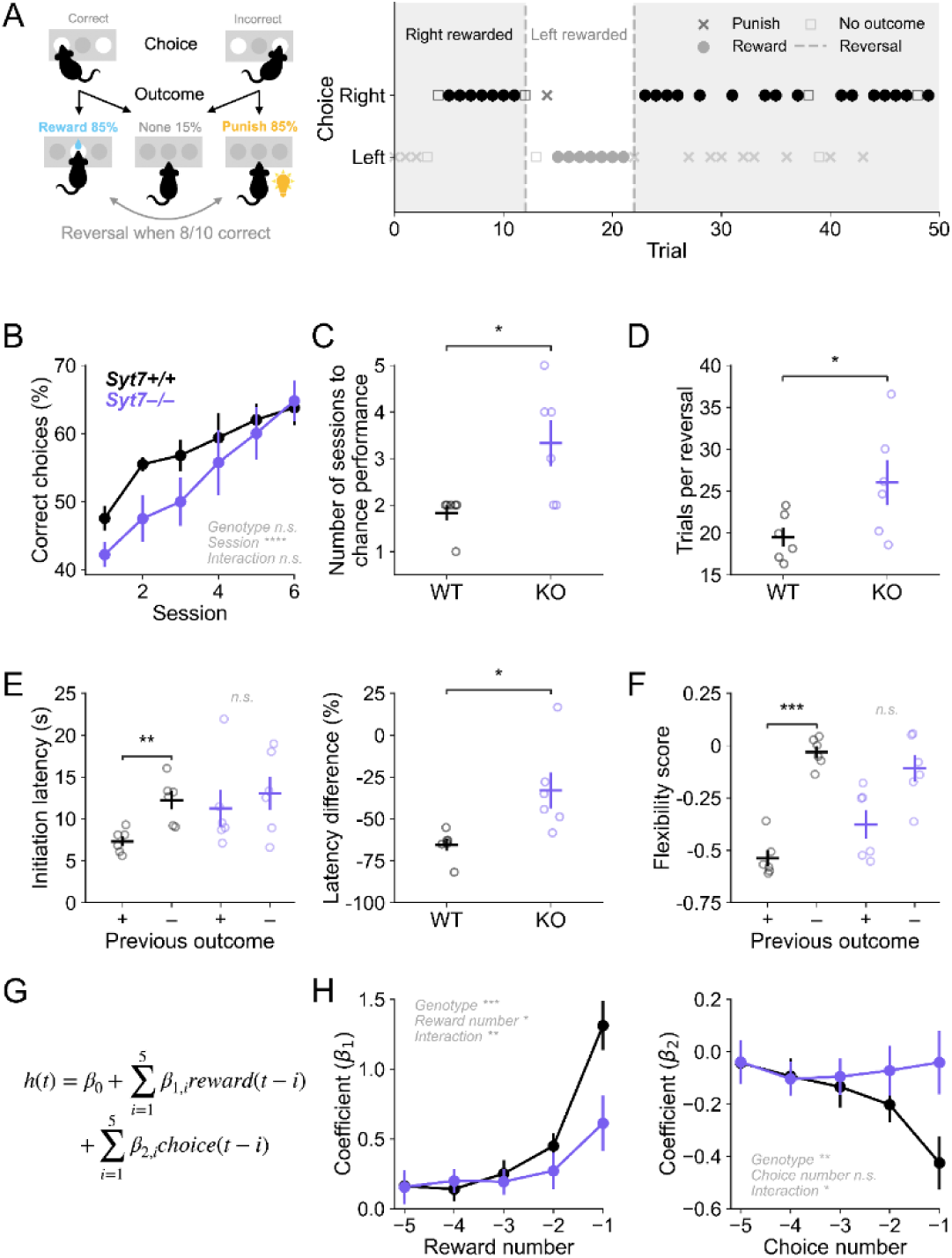
Knockout of Syt7 impairs outcome and trial history processing in probabilistic serial reversal learning. A) Left: Schematic of two-choice probabilistic serial reversal task. Right: Exemplar data showing choices, reward outcomes, and reversals for a WT mouse. B) Proportion of correct trials across training sessions for WT (N=6) and *Syt7* KO animals (N=6). C) Number of training sessions required to reach chance-level performance (50% of trials correct in a session). D) Number of trials between consecutive reversals, averaged across all sessions. E) Left: Latency to initiate trials after rewarded (+) and punished (–) trials, averaged across all sessions for each mouse. Right: Percentage difference in trial initiation latencies following rewarded trials compared to unrewarded trials. F) Flexibility scores calculated for choices made after rewarded (+) and punished (–) trials, calculated as difference between number of “shift” trials (choice not repeated) and number of “stay” trials (choice repeated) divided by total trials. G) Logistic regression model formula using outcomes and choices of preceding 5 trials to predict current trial choice. H) Fitted logistic regression model coefficients. Left: Influence of past five trial outcomes on current choice. Right: Influence of past five choices on current choice. Data points are expressed as mean ± SEM, with small hollow circles representing individual animal replicates. Statistical significance is shown as *p<0.05, **p<0.01, ***p<0.001, and ****p<0.0001.

To quantify how recent outcomes influenced choice behavior while controlling for differences in performance, we analyzed win-stay/lose-shift strategies. Flexibility scores were calculated for rewarded and punished trials as the difference between the proportions of “shift” choices and “stay” choices, such that more negative scores indicate more repeated choices and more positive scores indicate fewer repeated choices (Aarde et al., 2019). WT mice were significantly more likely to repeat rewarded choices than punished choices (*t*(5)=14.3, *p*=0.00010), whereas KO mice did now show a significant difference in shift/stay behavior by outcome (*t*(5)=3.1, *p*=0.073; Fig. 2F). Both groups exhibited similar near-random choice behavior following punished (WT -0.03±0.03; KO -0.11±0.06; *t*(10)=1.09, *p*=0.90) and no-outcome trials (WT -0.06±0.03; KO -0.10±0.03; *t*(10)=1.14, *p*=0.84), with no differences between punish and no-outcome conditions in either genotype (WT *t*(5)=1.0, *p*=0.36; KO *t*(5)=-0.05; *p*=0.96), indicating that the primary source of performance differences was in how each group responded to positive feedback from successful trials. Consistent with this, win-stay/lose-shift model fits trended toward higher lapse rates in KO mice (WT 0.38±0.01; KO 0.44±0.02; *t*(10)=-2.09, *p*=0.063), suggesting a greater tendency to make choices independent of recent reward history.

We further assessed history dependence using a logistic regression model to predict current choice from the five most recent choices and outcomes (Fig. 2G). WT choices were strongly influenced by prior choices and outcomes, particularly from the immediately preceding trial (Fig. 2H). In contrast, regression coefficients in KO mice were near zero across all 5 previous trials, indicating that recent choices (Main effect of genotype: β=-0.38±0.12, *z*=3.23, *p*=0.0013; Genotype × Choice number interaction: 0.090±0.04, *z*=2.54, *p*=0.011) and outcomes (Main effect of genotype: β=-0.66±0.17, *z*=2.21, *p*=0.00013; Genotype × Reward number interaction: -0.16±0.05, *z*=-3.11, *p*=0.0019) exerted a weaker influence on subsequent decisions.

Together, these results demonstrate that *Syt7* KO mice remain engaged and are capable of learning task structure, but their choices are less influenced by recent rewards, consistent with a deficit in integrating outcome history to update action values.

### Intact short-term memory and motor behaviors in Syt7 KO mice

Although the tasks described above did not include a delay period that required mice to remember the location of the sample port, the physical separation between ports required mice to use short-term memory to guide their movement. Thus, it is possible that KO mice performed more poorly due to memory deficits. This possibility is particularly noteworthy because a large number of computational studies have suggested that facilitation might serve as the substrate for information storage in short-term memory (Mongillo et al., 2008; Itskov et al., 2011; Hansel and Mato, 2013). To determine whether deficits observed in reversal tasks could be explained by impairments in short-term memory, we tested animals on multiple delay-dependent behavioral paradigms.

Across all tasks, KO mice performed comparably to WT controls. In the two-choice match-to-sample task with a 0-to 20-second delay between the sample and choice phases (Fig. 3A), accuracy improved across sessions (Main effect of session: *F*(20,400)=26.7, *p*<0.0001) and declined as a function of delay length for both genotypes (Main effect of session: *β*=-0.0008, *p*=0.02; effect of delay: *β*=-0.015, z=-5.67 *p*<0.0001; Fig. 3B,C). Although KO mice performed fewer trials in the shaping and delayed match phases of training (Extended Data 3-1A), there were no differences in choice or reward poke latencies indicative of altered motivation (Extended Data 3-1B).

**Figure 3:**
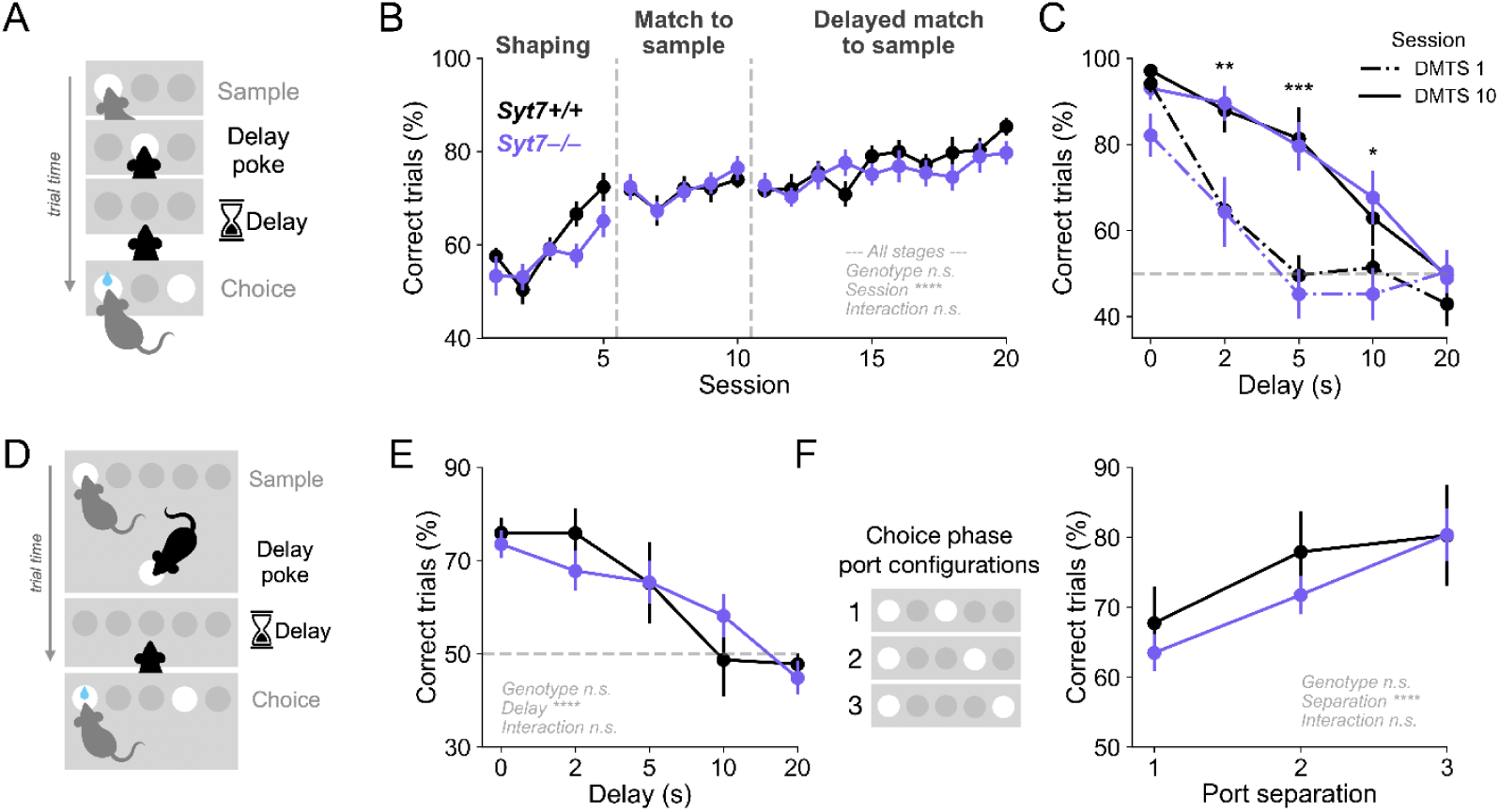
Short-term memory is unaffected by *Syt7* knockout. A) Schematic of two-choice delayed match-to-sample task. B) Proportion of correct trials across all sessions for WT (N=10) and *Syt7* KO animals (N=12). Dotted lines indicate transitions between training stages. C) Proportion of correct trials across delay lengths. Asterisks indicate statistically significant differences between delayed match-to-sample (DMTS) sessions 1 and 10 with genotypes pooled together. D) Schematic of five-choice delayed match-to-sample task. E) Proportion of correct trials across delay lengths for WT (N=3) and *Syt7* KO animals (N=6). F) Proportion of correct trials across different choice phase port configurations, each identified by the number of inactive ports between the two choice ports. Data is averaged from all trials with delay lengths less than 10 seconds. Data points are expressed as mean ± SEM, with small hollow circles representing individual animal replicates. Statistical significance is shown as *p<0.05, **p<0.01, ***p<0.001, and ****p<0.0001.

We further transitioned a subset of subjects to a five-choice version of the same task to increase difficulty (Fig. 3D). Once again, there was a strong relationship between delay length and choice accuracy, but this did not differ across genotypes (Main effect of delay: *F*(4,28)=20.5, *p*<0.0001; Main effect of genotype: *F*(1,7)=0.03, *p*=0.87; Fig. 3E). Performance also declined for both genotypes when the distance between the correct and incorrect choice ports was decreased (Main effect of port distance: *F*(2,14)=42.4, *p*<0.0001; Fig. 3F), indicating that the physical distance between illuminated choice ports affected task difficulty despite there being no change to the reward contingency.

To further assess spatial working memory, we tested WT and *Syt7* KO mice on an eight-arm radial arm maze, a canonical assay of memory-guided foraging behavior (Fig. 4A). Performance improved across training sessions (Main effect of session: *β*=-1.0, *p*<0.0001; Fig 4B) as animals in both groups adopted a sequential turning strategy, making more 45° turns (Main effect of session: *F*(2,20)=48.7, p<0.0001; Fig 4C, left). However, performance was disrupted by the addition of an enforced delay period between consecutive arm entries. KO mice made significantly more errors than WT mice on the first day of confinement (*t*(5)=4.96, *p*=0.0023; Fig. 4B). This was partially driven by an increase in total entries in KOs compared to WTs (*t*(10)=-5.49, *p*=0.0003; Fig 4D), but with KOs exhibiting a reduced proportion of correct entries (*t*(10)=3.4, *p*=0.0067; Fig. 4E). Both genotypes showed a similar proportion of 45° turns to the first day of unconfined exploration, demonstrating that confinement inhibited use the sequential turning strategy (Unconfined day 1 vs confined day 1: WT *t*(5)=1.3, *p*=0.8, KO *t*(5)=0.08, *p*=1.0).

**Figure 4:**
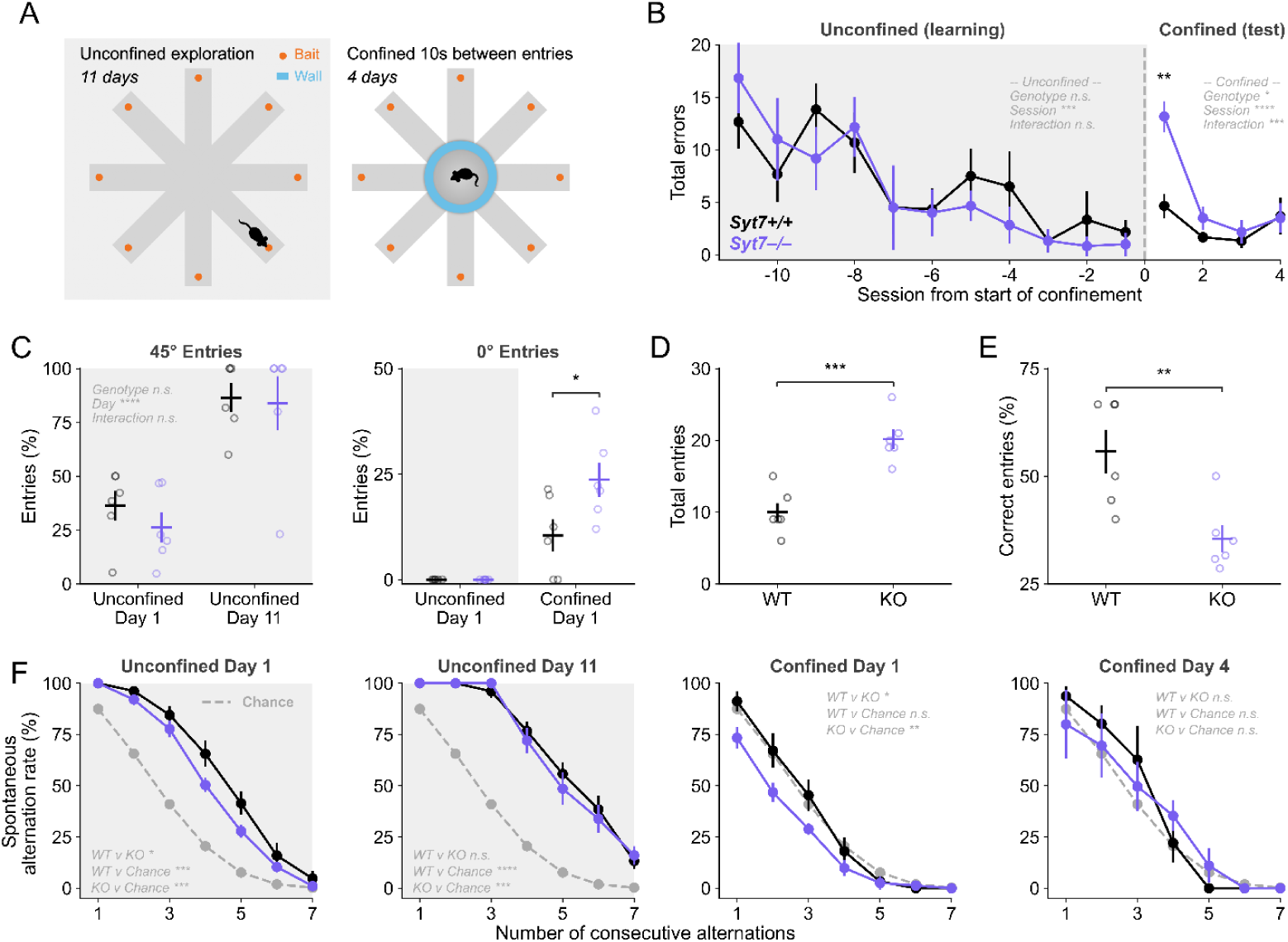
Spatial strategy learning is normal but transiently disrupted by delay in *Syt7* KO. A) Schematic of radial arm maze task. B) Number of entries into previously visited (no longer rewarded) arms across training sessions for WT (N=6) and *Syt7* KO animals (N=6). C) Left: Proportion of entries made into the immediately adjacent arms on day 1 of unconfined exploration and day 11 of unconfined exploration (e.g. exiting arm 3 and entering arm 2 or arm 4). Left: Proportion of entries made into the most recently visited arm on day 1 of unconfined learning and day 1 of confined testing (e.g. exiting arm 3 and re-entering arm 3). D) Total arm entries in the first session with confinement. E) Total baited arm entries in the first session with confinement. F) Tendency to spontaneously alternate (consecutively visit unique arms) across selected sessions, e.g. entering a new arm after the preceding entry is 1 consecutive alternation, while entering each of the 8 arms without repeat is 7 consecutive alternations. Dotted line indicates alternation rate expected by random chance and statistical annotations reflect comparisons of area under the curve. Data points are expressed as mean ± SEM, with small hollow circles representing individual animal replicates. Statistical significance is shown as *p<0.05, **p<0.01, ***p<0.001, and ****p<0.0001.

The delay period confinement substantially disrupted spatial navigation, as mice showed an increase in 0° entries (returning to the arm just visited), despite neither genotype exhibiting this behavior during non-confined exploration (Fig. 4C, right). To better understand this behavior, we analyzed performance by determining the spontaneous alternation rate for every possible number of consecutive alternations (1 through 7) and calculating the area under this curve. Both early and late in unconfined exploration, both genotypes showed alternation rates significantly above chance (Fig. 4F, left), with KOs alternating less than WTs on the first session (*t*(10)=2.35, *p*=0.04). The introduction of a delay between consecutive entries decreased alternation rates to chance level, with KOs alternating significantly less what would be expected by chance on the first day of confinement (*t*(10)=14.6, *p*<0.0001; Fig. 4F, right). Notably, this impairment was transient, with KO performance recovering across subsequent sessions, suggesting a deficit in adaptive behavior following abrupt changes in task demands, rather than a persistent impairment in spatial memory itself.

To again rule out the possibility of motor deficits, we tested mice in a rotarod assay and found the average latency to fall did not significantly differ between genotypes (Main effect of genotype: *F*(1,16)=2.23, *p*=0.15; Fig. 5A). In an open field, we observed no differences in locomotion or anxiety-like behavior, as measured by total distance traveled (*U*=19, *p*=0.94) and time spent in the center of the arena (*t*(10)=0.044, *p*=0.97; Fig. 5B). We further examined behavior in a Y-maze to assess spontaneous alternation as a measure of short-term memory in the absence of reward-motivated exploration (Fig. 5C). WT and KO mice demonstrated similar distance traveled (*t*(7)=0.24, *p*=0.82), arm entries (*t*(7)=0.67, *p*=0.52), and spontaneous alternation rates (*t*(7)=-0.94, *p*=0.38), suggesting intact memory of recently visited arms (Fig. 5D).

**Figure 5:**
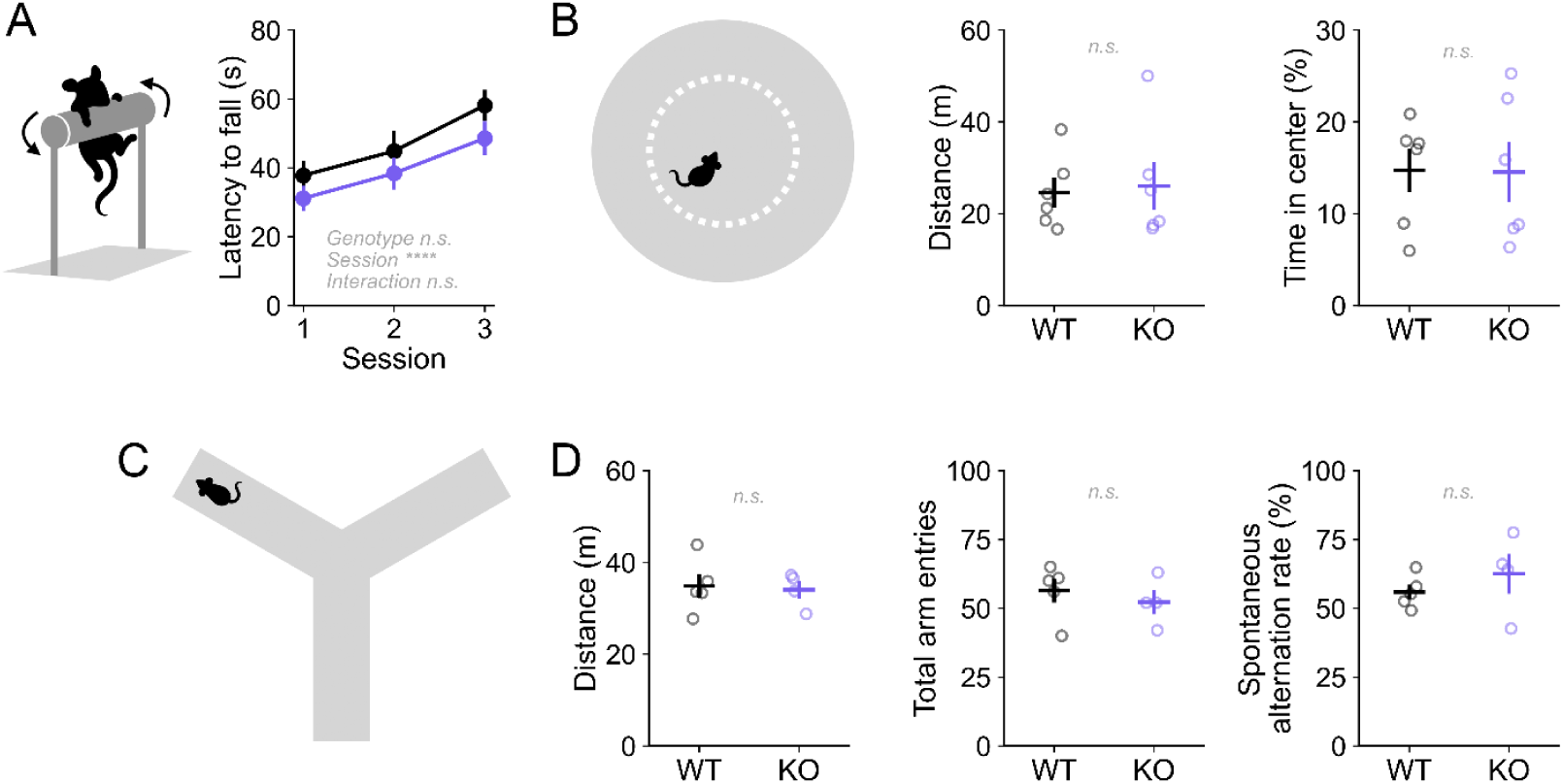
Motor behavior is normal in *Syt7* KO. A) Average latency to fall from rotating rod across 3 test consecutive days for WT (N=9) and *Syt7* KO animals (N=9). B) Schematic (left), distance traveled (center), and time in center third of open field (right) for WT (N=6) and *Syt7* KO animals (N=6). C) Y maze task schematic. D) Distance traveled (left), total arm entries (center), and spontaneous alternation rate (right) of Y maze exploration for WT (N=5) and *Syt7* KO animals (N=4). Data points are expressed as mean ± SEM, with small hollow circles representing individual animal replicates. Statistical significance is shown as *p<0.05, **p<0.01, ***p<0.001, and ****p<0.0001.

Together, these findings demonstrate that short-term spatial memory and memory-guided navigation are preserved in *Syt7* KO mice. The behavioral impairments observed in reversal paradigms are therefore unlikely to reflect memory limitations and instead point to a more specific deficit in updating behavior based on changing task contingencies. This is consistent with the disrupted navigation behavior seen in KOs following the change in task structure.

## Discussion

The ability to flexibly adapt behavior to changing environmental demands is a defining feature of cognition. Cognitive flexibility requires a balance of stability and adaptability: animals must maintain learned representations when the environment is stable, yet rapidly update behavior when contingencies change. Although decades of work have identified synaptic plasticity as a fundamental substrate of neural computation, the specific cognitive functions supported by distinct forms of plasticity remain incompletely understood. Among these mechanisms, short-term synaptic facilitation has occupied a particularly compelling position. Since its earliest descriptions nearly a century ago, facilitation has been proposed to transiently retain information, shape temporal integration, and support flexible computations across neural circuits (Feng, 1940). Yet despite extensive theoretical and physiological study, directly linking synaptic facilitation to behavior has remained challenging. The identification of SYT7 as a key driver of facilitation makes it possible to examine how selective disruption of this form of plasticity alters adaptive behavior *in vivo*.

Our results suggest that loss of SYT7-driven facilitation disrupts how recent outcomes influence future choices. In a probabilistic environment, KO mice showed reduced outcome-dependence in both choices and initiation latencies, indicating that recent rewards had less impact on behavior. In a deterministic rule-reversal task, this manifested as increased perseverative errors, consistent with difficulty adapting behavior to new task demands. Across tasks, these effects point to a reduced influence of recent outcomes on behavior rather than a global impairment in learning.

## Synaptic computations

At a computational level, flexibility requires assigning appropriate weight to recent experience. If past information is weighted too heavily, behavior becomes perseverative; if recent information is weighted too lightly, adaptation is slowed. Synaptic facilitation is well positioned to implement such recency weighting because its efficacy depends on immediately preceding activity. Although the intrinsic timescale of facilitation is on the order of hundreds of milliseconds, its effects on the input-output relationship between presynaptic firing rate and postsynaptic response can propagate through downstream networks on the order of seconds to tens of seconds (Dao Duc et al., 2015; Seeholzer et al., 2019).

Because facilitating synapses transiently amplify recently active pathways, they can bias network-wide activity toward particular states and expand the repertoire of accessible activity regimes (Barri and Mongillo, 2022). The seminal work of Tsodyks et al. (1998) formalized this intuition in recurrent network models, demonstrating that short-term plasticity fundamentally shapes the computational capacity of neural circuits. Our findings connect this theoretical foundation to a specific behavioral function.

Within reinforcement-learning frameworks, the present results could be interpreted as a reduction in effective learning from recent outcomes, or equivalently, as a shortening of the temporal window over which outcomes influence value estimates. Whether SYT7 loss primarily reduces learning rate, impairs forgetting, or destabilizes action-outcome representations remains an open question. Future computational modeling will be essential to distinguish among these accounts and to relate the timescales of synaptic facilitation to the behavioral dynamics observed here.

At the synaptic level, SYT7 supports short-term facilitation and sustained neurotransmitter release (Jackman et al., 2016; Weingarten et al., 2024). These properties could both amplify or prolong the impact of recent activity within task-relevant circuits. One interpretation is that this mechanism extends the temporal window over which outcomes influence decision-making, allowing recent rewards to more strongly bias subsequent choices. In the absence of SYT7, neuronal population coding for recent outcomes may decay more rapidly or exert weaker influence, leading to diminished trial-by-trial updating.

A key constraint on this interpretation is that Syt7 KO mice acquired initial task contingencies normally. This indicates that SYT7 is not required for forming stable action-outcome associations over longer timescales, but instead becomes particularly important for adjusting behavior based on recent experience. Reversal learning places a greater demand on this process, as previously learned associations must be overridden and replaced using ongoing feedback. Under these conditions, reduced sensitivity to recent outcomes would be expected to selectively impair performance, as observed here.

An alternative, not mutually exclusive, possibility is that SYT7 supports the updating of task representations required for flexible behavior. In this view, intact acquisition may be supported by repeated, consistent reinforcement, whereas reversal requires both suppressing previously reinforced responses and generalizing a new rule across trial types (Izquierdo et al., 2017). The elevated perseverative errors in KO mice are consistent with difficulty disengaging from previously learned contingencies following a change in reinforcement. Such a deficit could arise from weakened outcome, impaired inhibitory control, or decreased flexibility in updating established behavioral strategies.

Distinguishing among these possibilities will require future work.

Task structure may further modulate these demands. Prior work suggests that match-to-nonmatch and nonmatch-to-match reversals are not behaviorally equivalent, potentially due to differences in the strategies required to solve them (Dunnett et al., 1989; Yhnell et al., 2016). The match-to-nonmatch reversal used here can be conceptualized as a conditional discrimination reversal, in which the sample port defines the correct response. Such tasks engage medial prefrontal cortex (Shaw et al., 2013) and place strong demands on integrating recent outcomes with conditional rules – a context in which SYT7-dependent processes may be particularly important.

More broadly, computational and *in vitro* work demonstrate that short-term plasticity shapes how synapses respond to patterned activity, with facilitating synapses preferentially transmitting high-frequency or burst inputs (Zucker and Regehr, 2002). In circuits with task-related dynamics, such as phasic responses in frontal cortex or burst-pause activity in striatal interneurons (Bissonette et al., 2015; Zucca et al., 2018; Ma et al., 2022), short-term plasticity may selectively enhance behaviorally relevant signals. Disruption of facilitation could therefore alter how outcome-related activity propagates through frontostriatal circuits, which are known to support reversal learning and are disrupted in models showing impaired flexibility with intact acquisition (Kehagia et al., 2010).

An additional interpretation arises from the established role of SYT7 in dopaminergic signaling. Beyond its function at conventional glutamatergic synapses, SYT7 regulates short-term facilitation and sustained dopamine release, particularly during phasic activity patterns (Hikima et al., 2022; Lebowitz et al., 2024). Dopamine neurons encode reward prediction errors that drive updating of action values and behavioral policies (Schultz, 2019). Within this framework, the reduced win-stay/lose-shift distinction, diminished history dependence, elevated perseverative errors, and attenuated outcome-dependent changes in initiation speed observed in KO mice could all reflect weakened transmission of recent reward information through dopamine-dependent learning mechanisms. This interpretation is consistent with intact initial acquisition alongside impaired adaptation following contingency changes, suggesting a selective deficit in outcome-guided updating rather than a global learning impairment (Izquierdo et al., 2017). Although our experiments do not directly assess dopamine signaling, the convergence between the known synaptic functions of SYT7 and the behavioral phenotype observed here makes altered dopaminergic reinforcement signaling an important hypothesis for future investigation.

## Future directions

Our results add behavioral evidence to a literature that has largely characterized facilitation at the cellular and circuit levels. Theoretical work has long proposed that facilitating synapses are particularly suited to transmitting information about recent activity history, effectively functioning as short-term memory buffers within neural circuits (Zucker and Regehr, 2002; Abbott and Regehr, 2004). *In vitro* studies have demonstrated that facilitation shapes the gain and timing of synaptic transmission in a use-dependent manner, enabling circuits to differentially weight inputs based on their recent firing history.

Our findings suggest that this biophysical property has functional consequences at the behavioral level, contributing specifically to the recency-weighted updating that underlies adaptive decision-making.

An important next step will be to embed the present findings within a formal computational framework. Computational models of reinforcement learning that incorporate synaptic facilitation as a mechanism for recency weighting could, in principle, reproduce the trial-by-trial behavioral signatures observed here – including the reduced history dependence of choice and latency and the pattern of perseverative errors during reversal. Such models would also generate testable predictions about the effects of SYT7 loss under varying task parameters, including reward probability, reversal frequency, and trial spacing. More mechanistic models, grounded in the network-level consequences of SYT7-dependent facilitation (Dao Duc et al., 2015; Seeholzer et al., 2019), could further bridge the gap between rapid synaptic dynamics and the behavioral timescales observed here. Together, computational and experimental approaches will be essential for understanding how short-term plasticity shapes the circuit computations that support flexible behavior.

Importantly, several alternative explanations are not supported by our data. KO mice showed intact performance on delay-based tasks and in the radial arm maze, indicating preserved short-term spatial memory. This is noteworthy given the substantial body of theoretical work proposing that facilitation supports information maintenance for short-term memory (Abbott and Regehr, 2004; Mongillo et al., 2008), and suggests that the behavioral relevance of facilitation may be more specific or context-dependent than these frameworks predict. Measures of task engagement and reward retrieval were also comparable between genotypes, arguing against motivational or motor confounds. Together, these findings support a relatively selective impairment in outcome-guided behavioral updating.

Several limitations should be considered. Our experiments do not localize the circuit mechanisms through which SYT7 acts, and in the future conditional *Syt7* depletion from specific circuits or neuronal populations will be required to identify the relevant brain regions and cell types. While our behavioral analyses support a deficit in outcome-dependent updating, computational modeling could help distinguish between reduced outcome sensitivity, impaired inhibitory control, and changes in representational stability. Finally, it remains to be determined how broadly these findings generalize across tasks and behavioral domains.

In summary, our results suggest that SYT7-driven facilitation plays a key role when behavior must be updated based on recent experience. These findings establish a direct link between a molecular mediator of synaptic short-term plasticity and higher-level cognitive function. More broadly, they suggest that short-term facilitation may contribute to a fundamental computational challenge faced by adaptive organisms: balancing the stability of established knowledge against the need to rapidly incorporate new information. While long-term plasticity is well suited to support durable learning, short-term facilitation may provide a complementary mechanism that transiently amplifies representations of recent outcomes precisely when actions must change. By connecting the molecular machinery of facilitation to outcome-guided behavioral updating, the present work suggests that short-term synaptic plasticity is not merely a biophysical property of individual synapses, but a mechanism with measurable and specific consequences for how the brain navigates a changing world.

## Author Contributions

CLL, AMB, and SLJ designed research; CLL, AMB, MK, JG, and AMA performed research; CLL analyzed data; CLL and SLJ wrote the paper.

## Acknowledgements

Research reported in this publication was supported by the Whitehall Foundation, the Medical Research Foundation of Oregon, and the National Institute of Neurological Disorders and Stroke of the National Institutes of Health under Award Numbers R01NS135048 (SLJ) and F31NS141376 (CLL). The content is solely the responsibility of the authors and does not necessarily represent the official views of the National Institutes of Health. We thank Suzanne H. Mitchell, Atheir I. Abbas, Alex Sonneborn and members of the Jackman lab for comments on the manuscript. Generative AI (ChatGPT 5.5 by OpenAI) was used in the preparation of this manuscript for polishing language and improving word count.

## Multimedia, Figures, and Tables

## Statistical tables

Fig 1

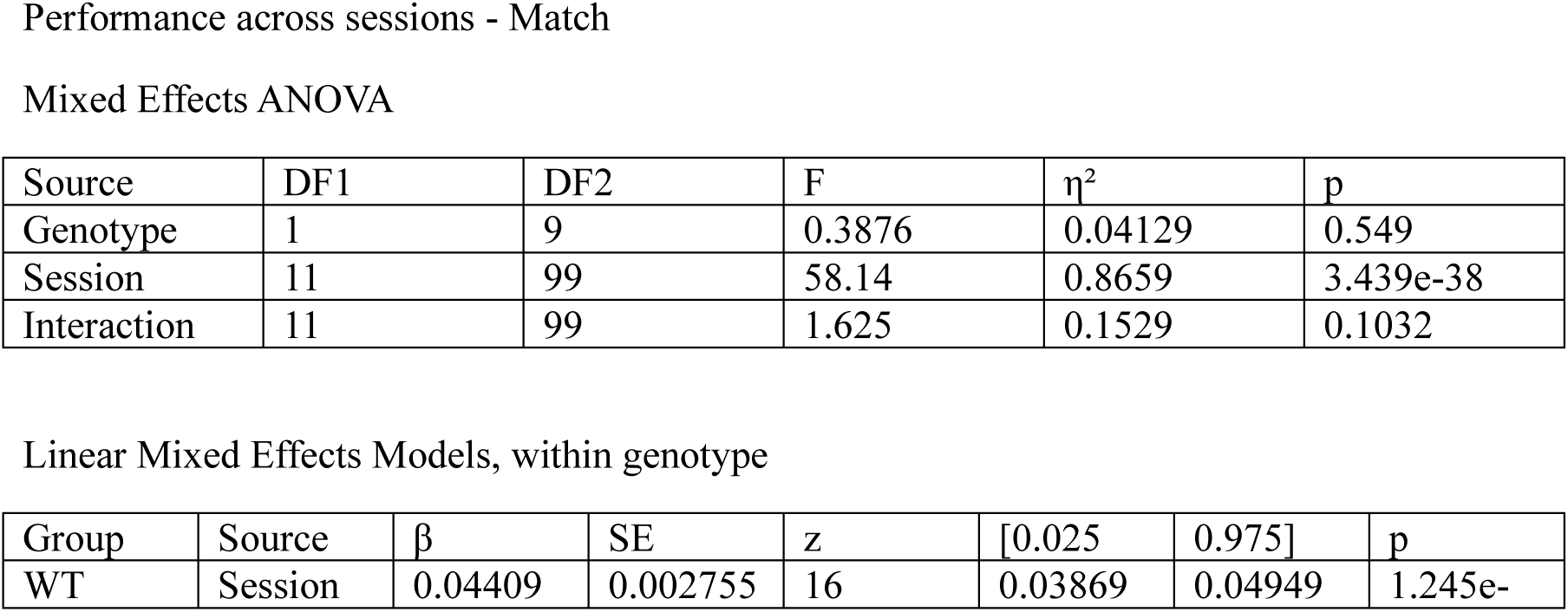
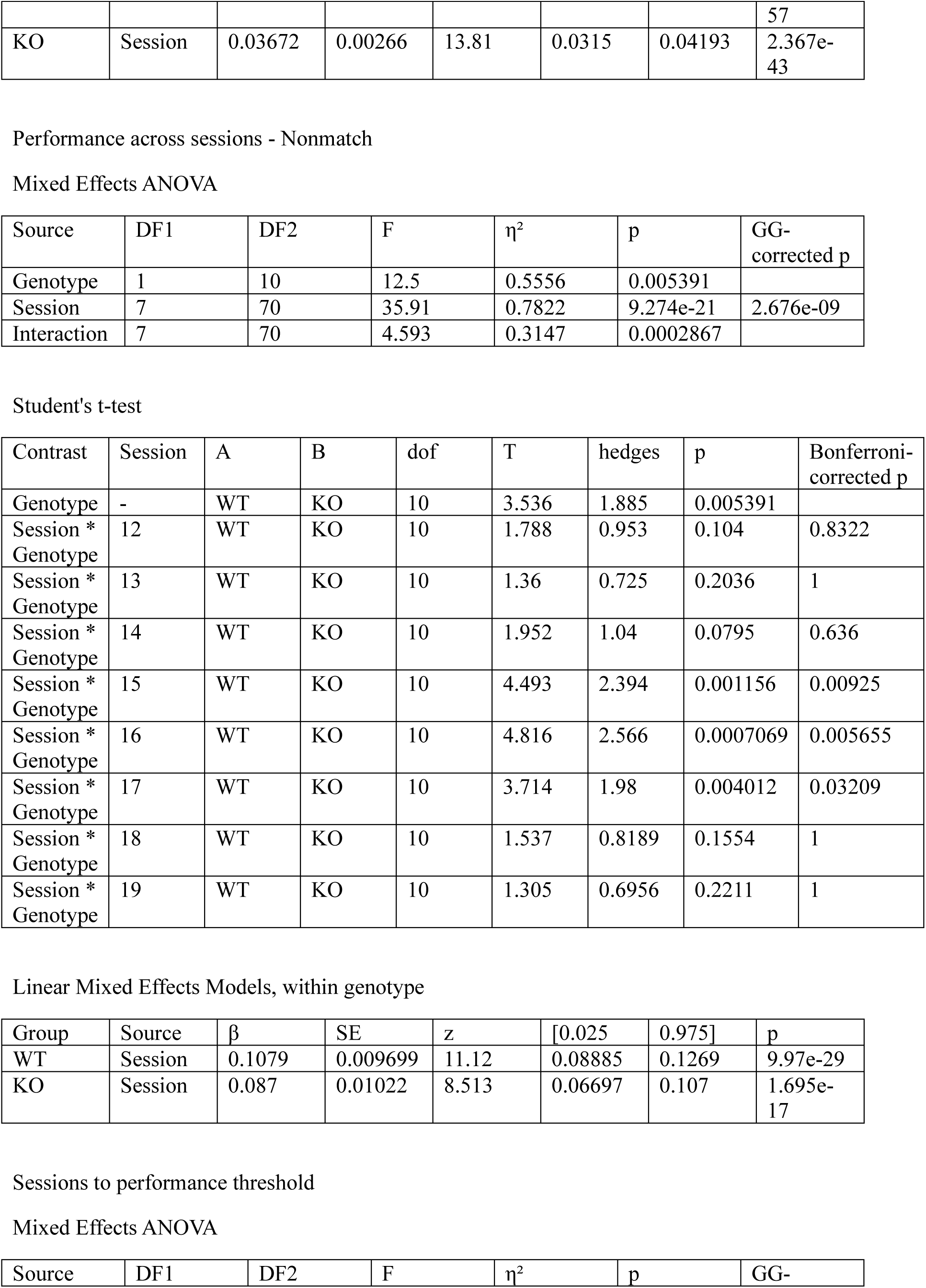
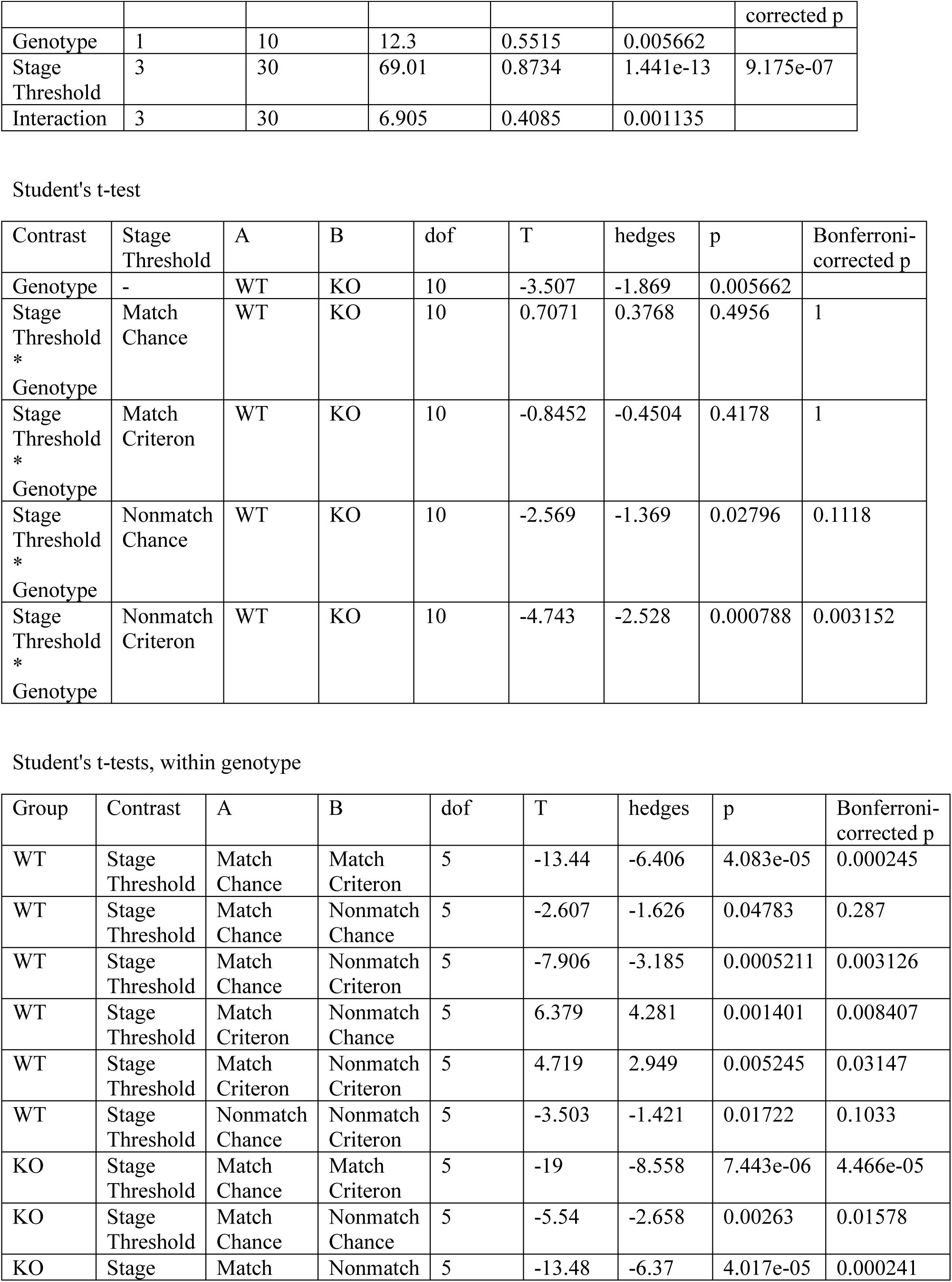
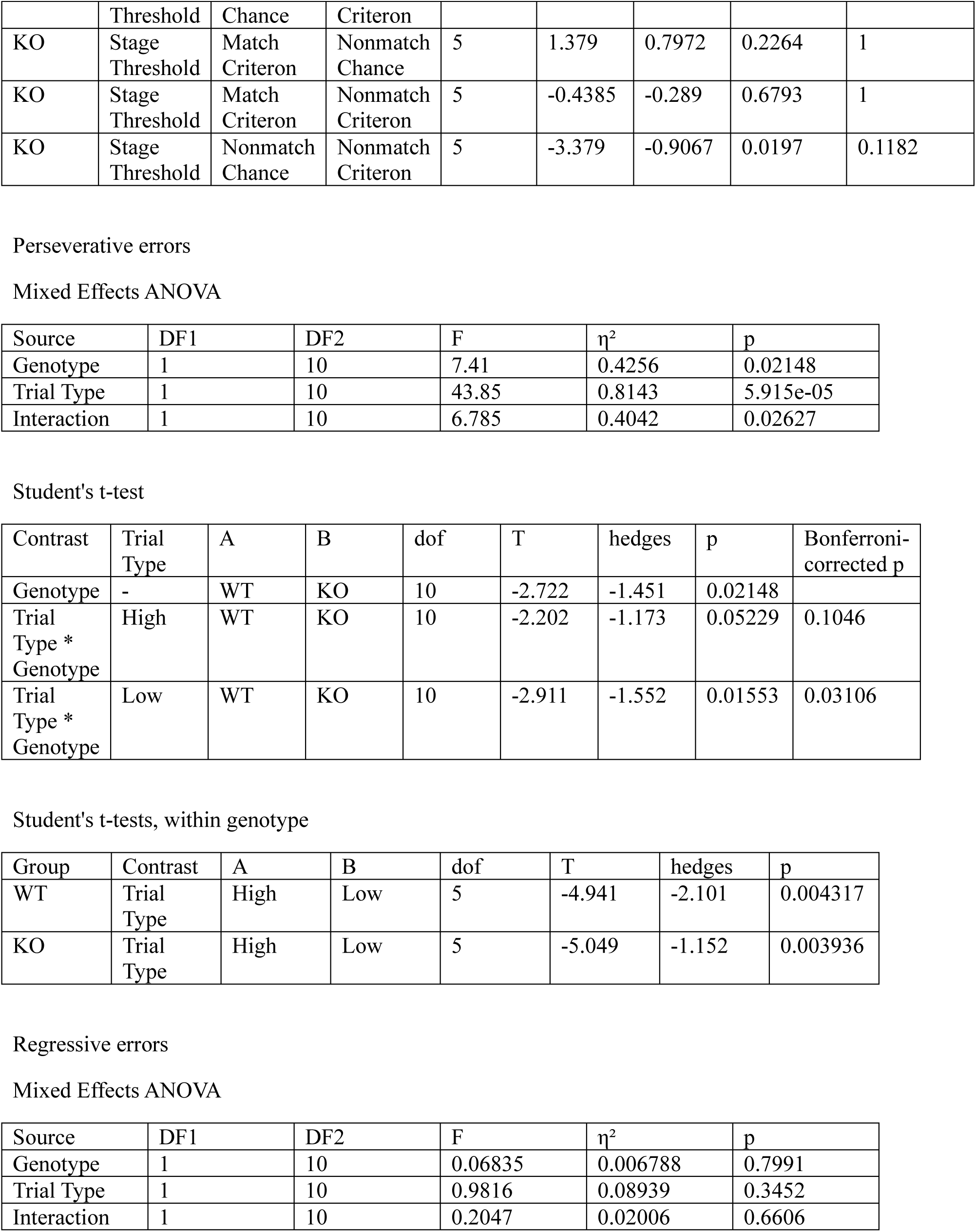
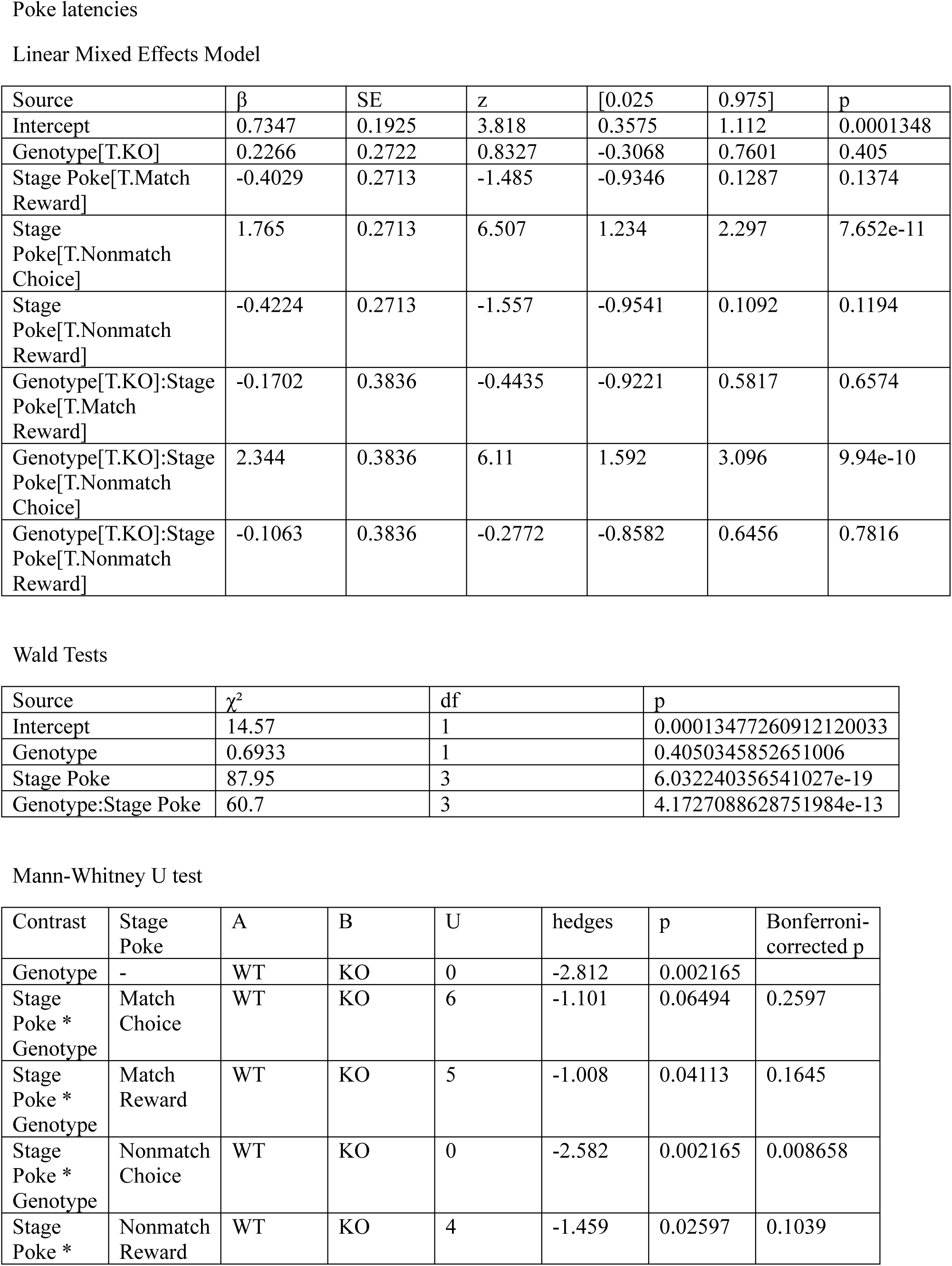
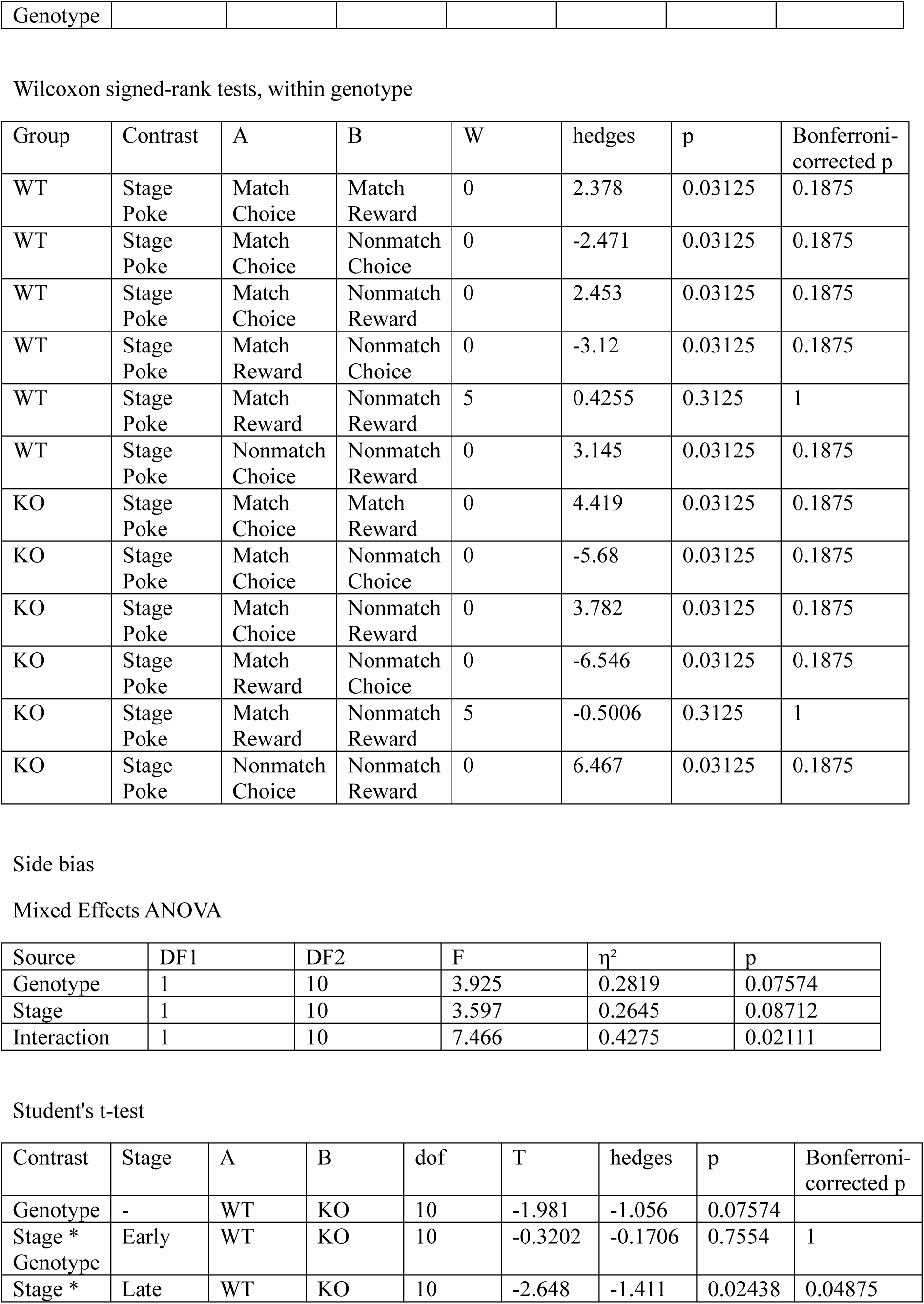
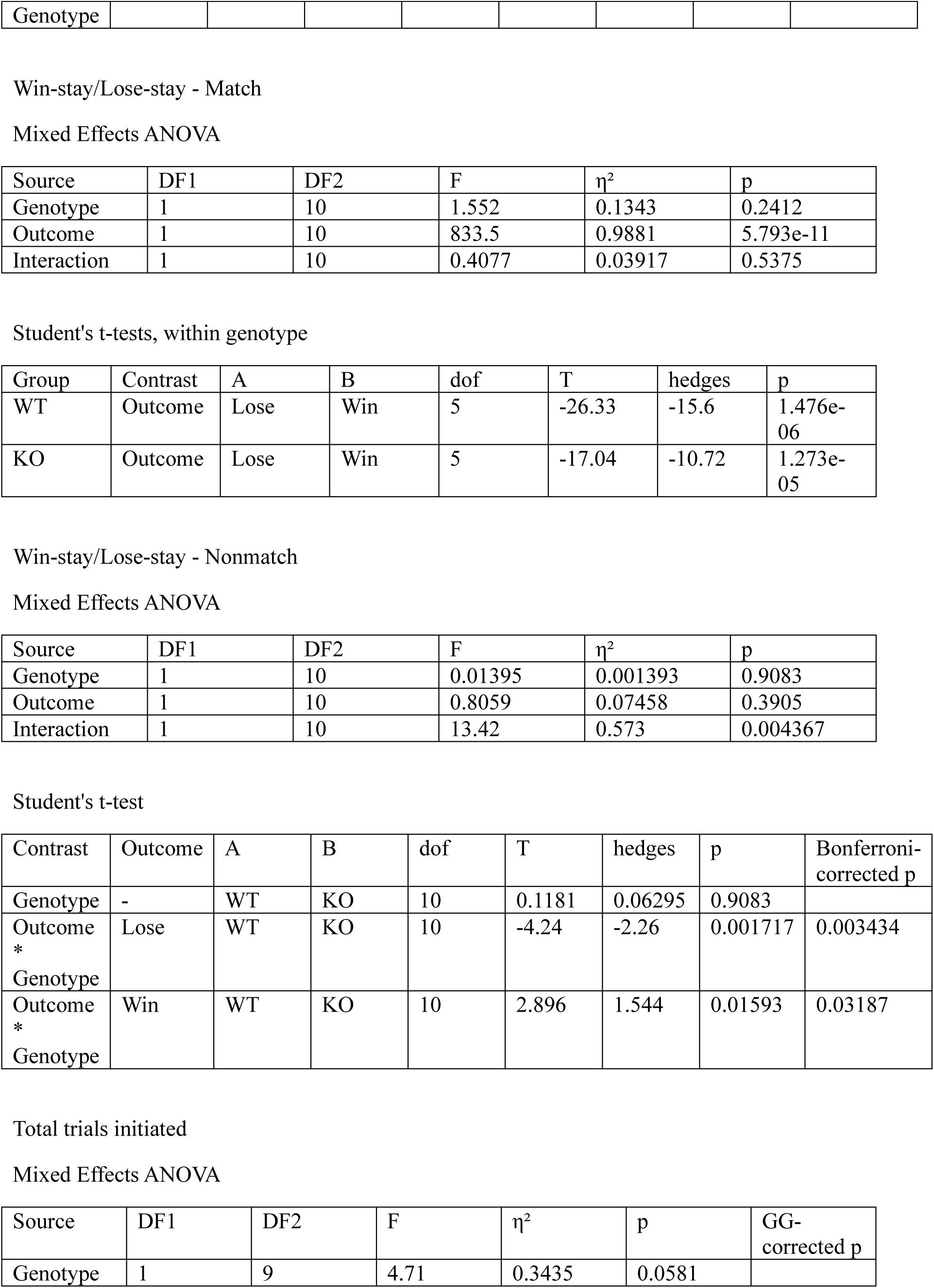
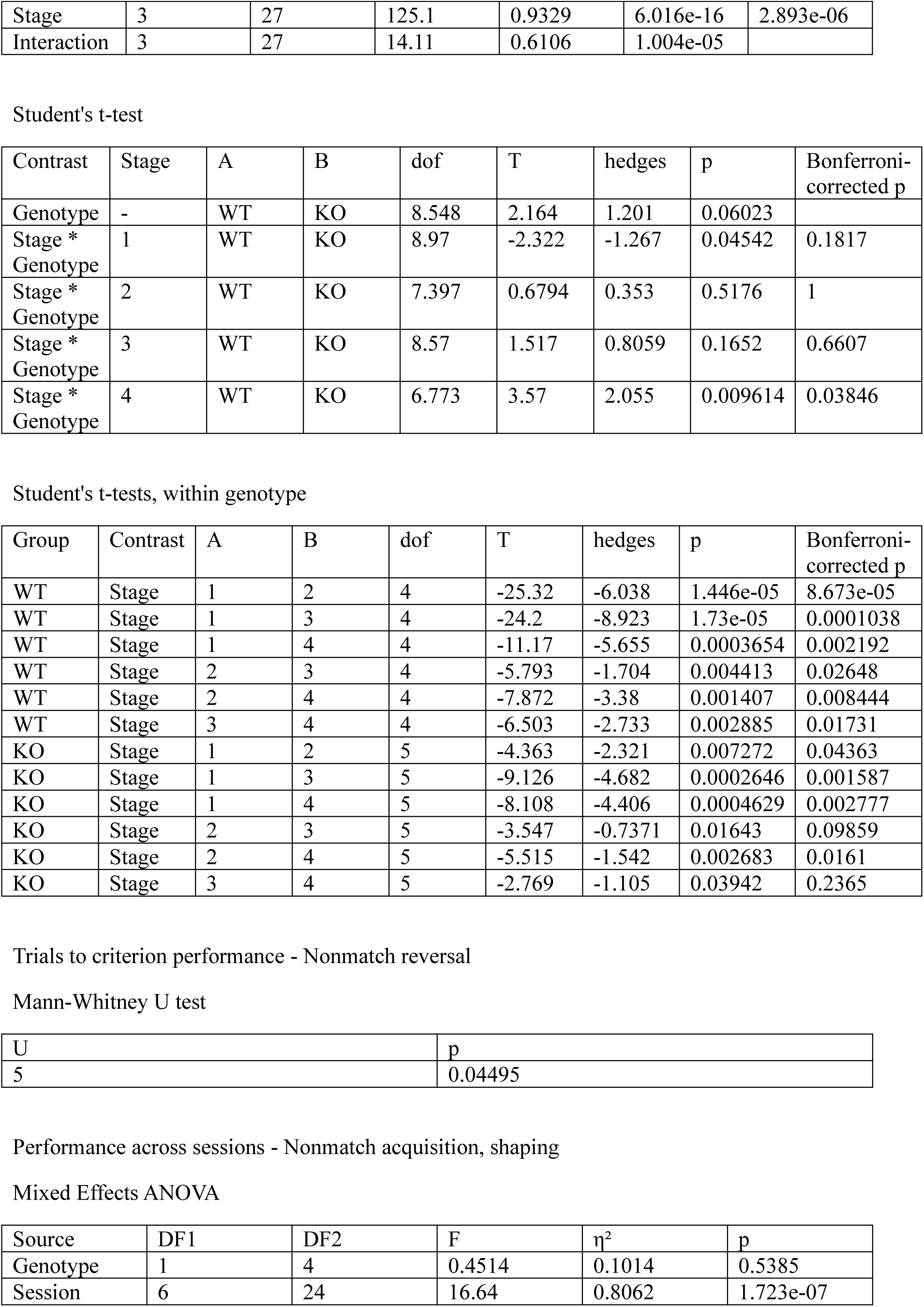
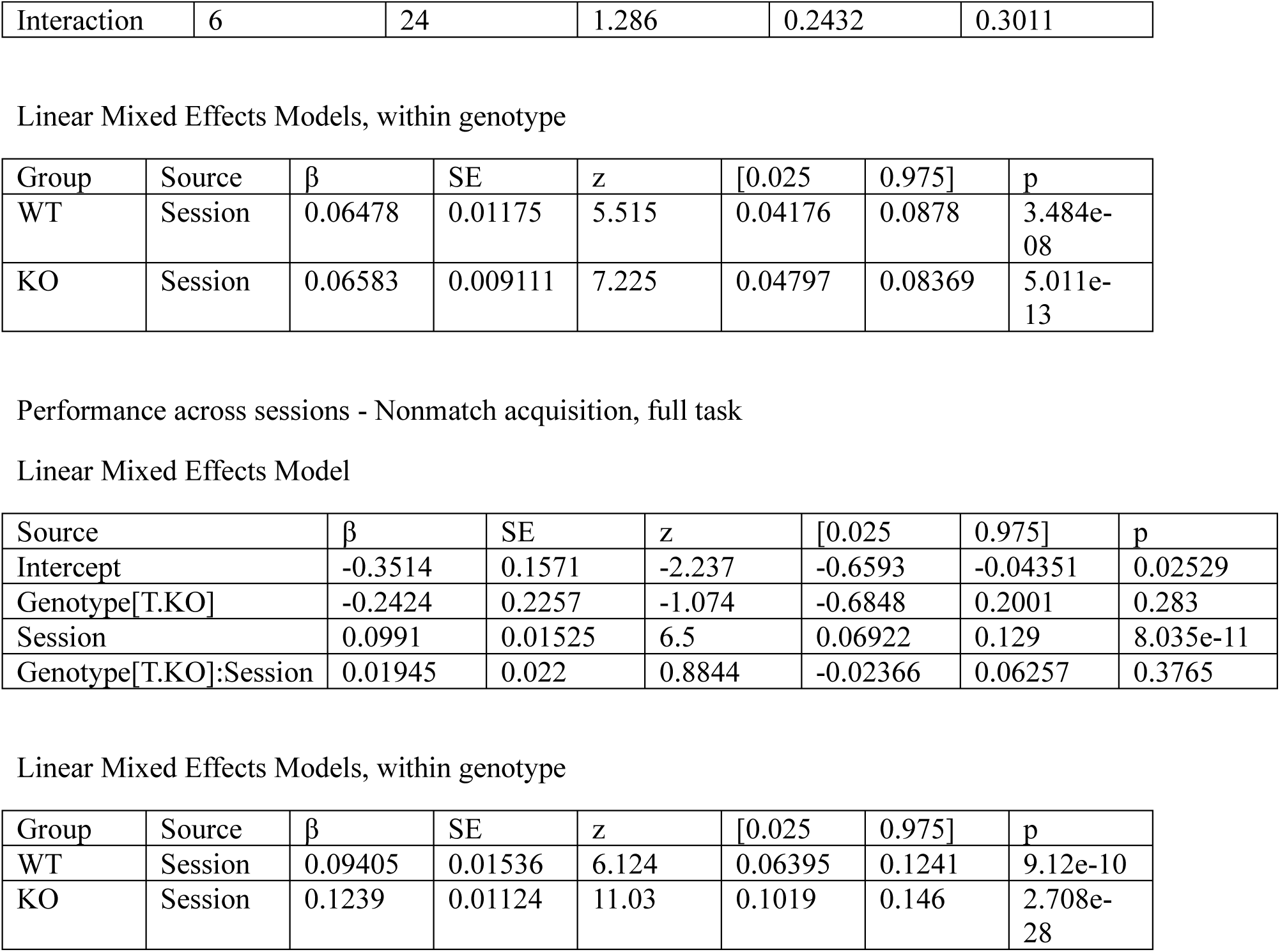

Fig 2

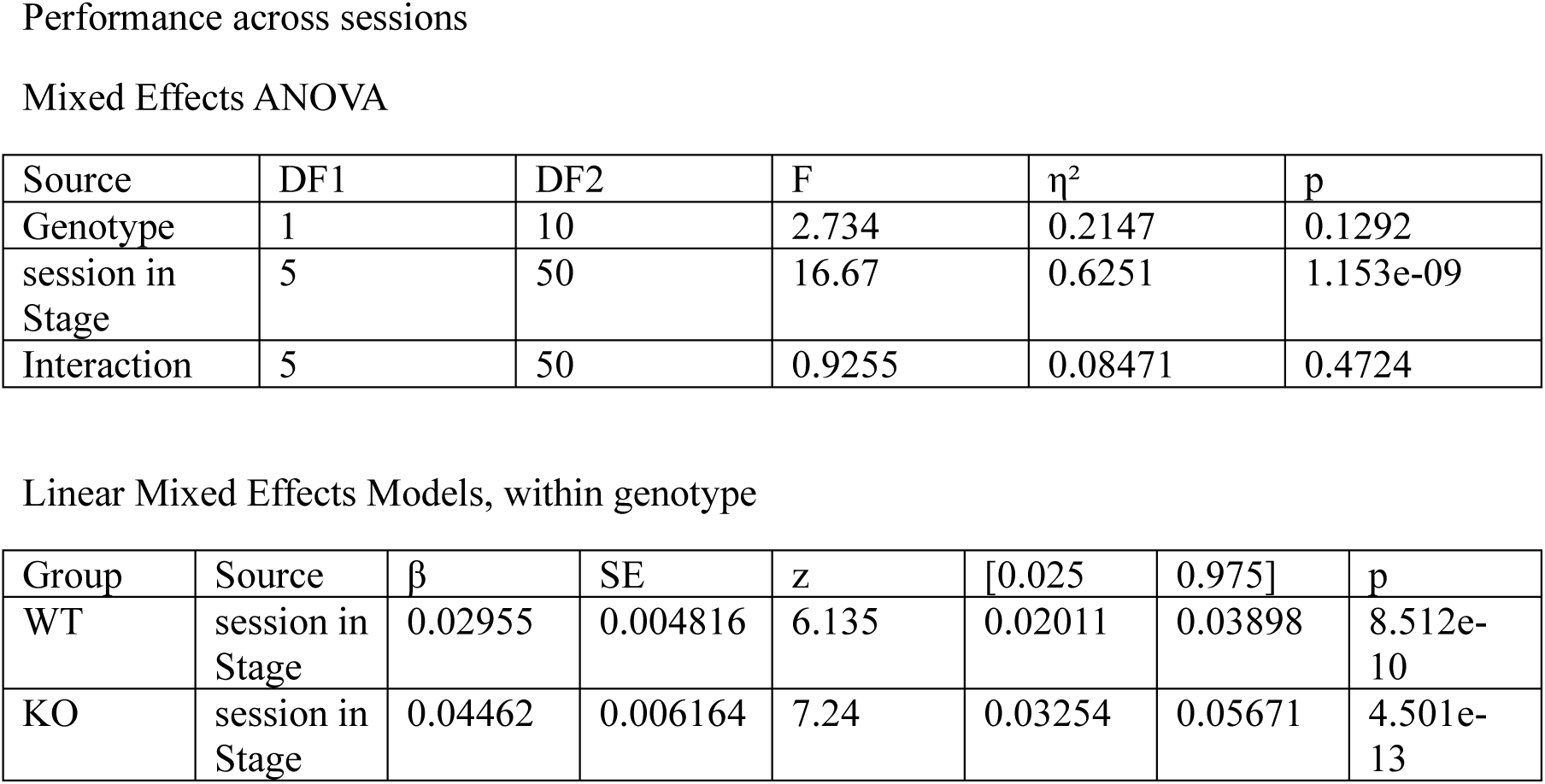
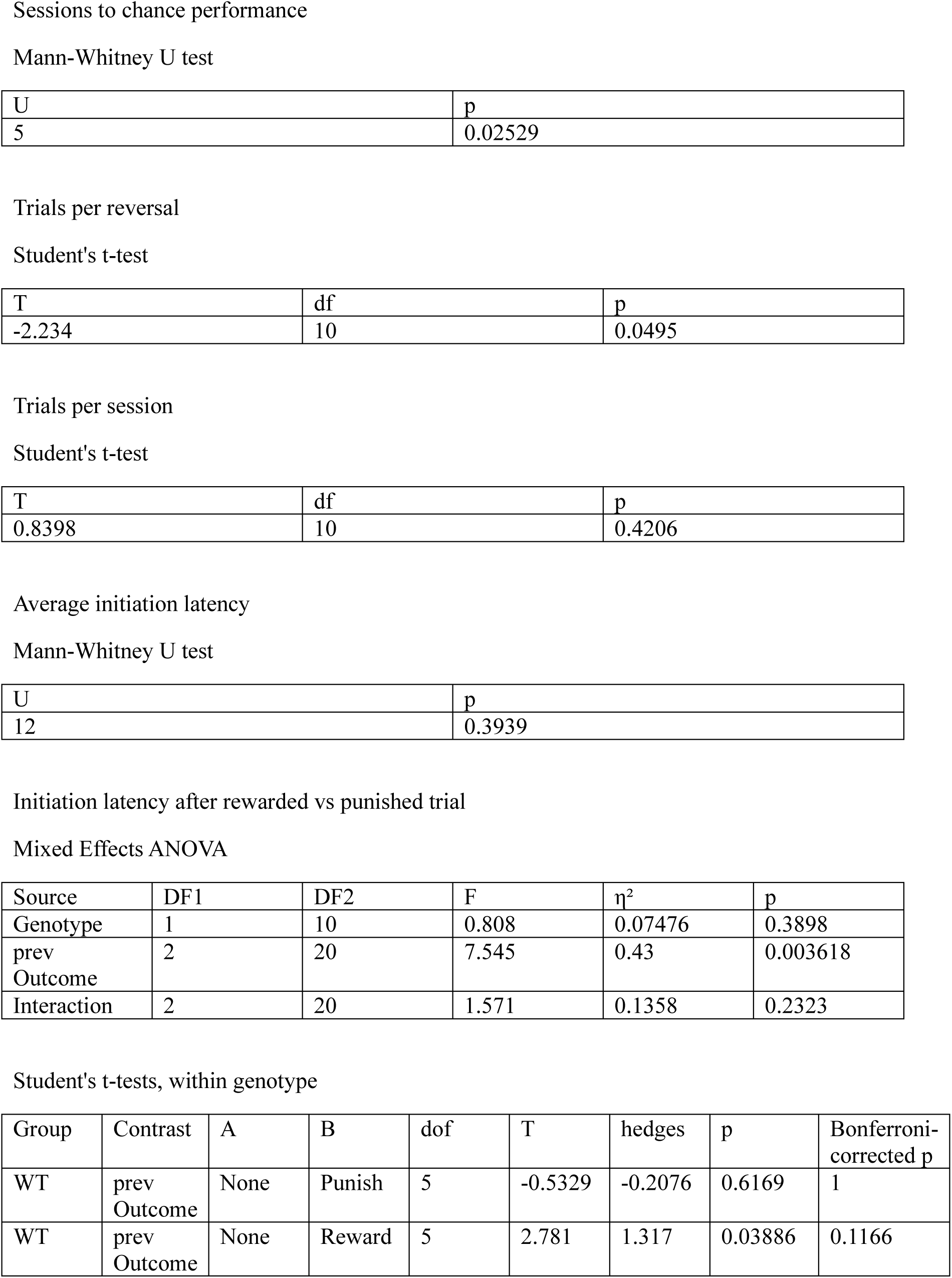
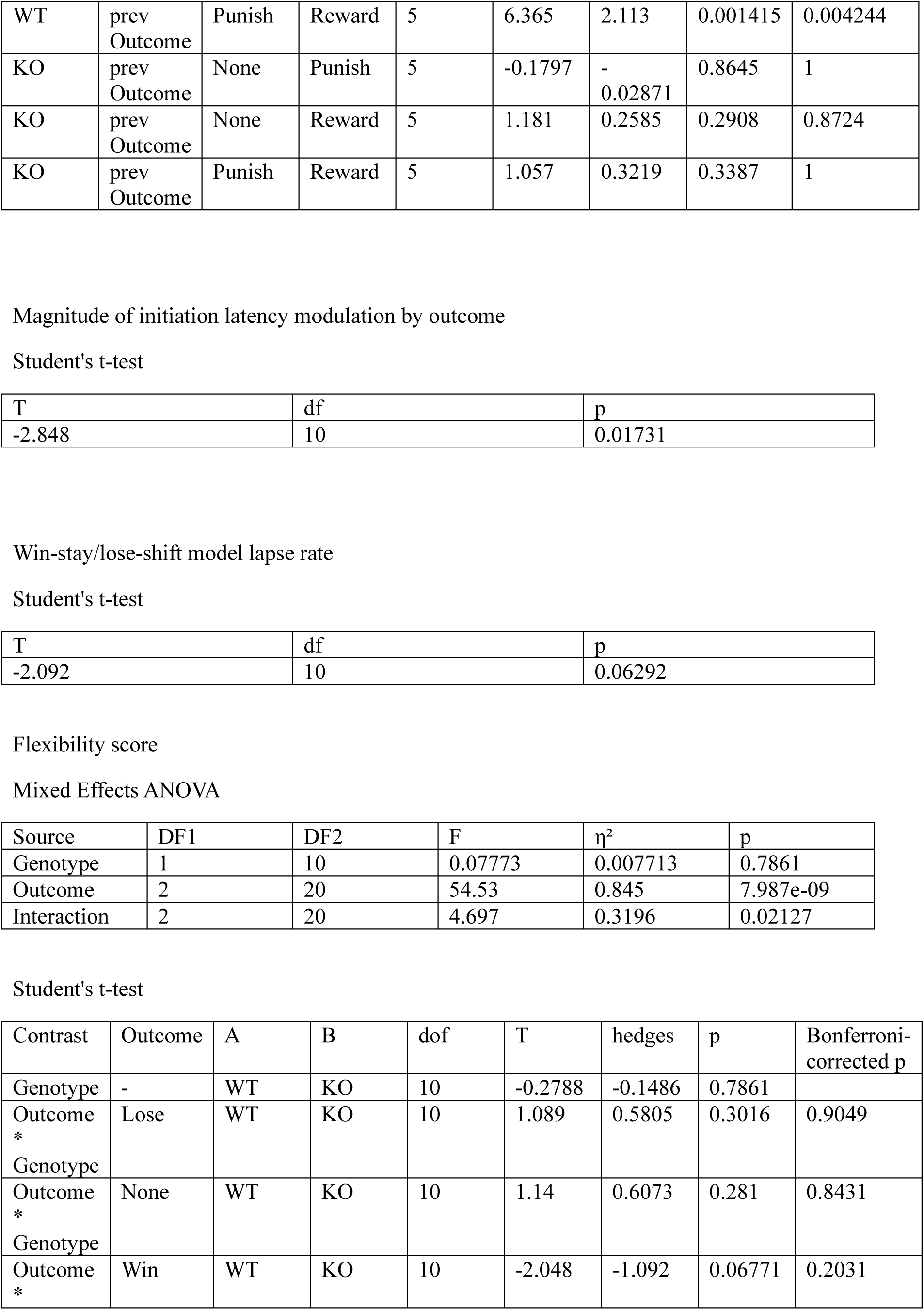
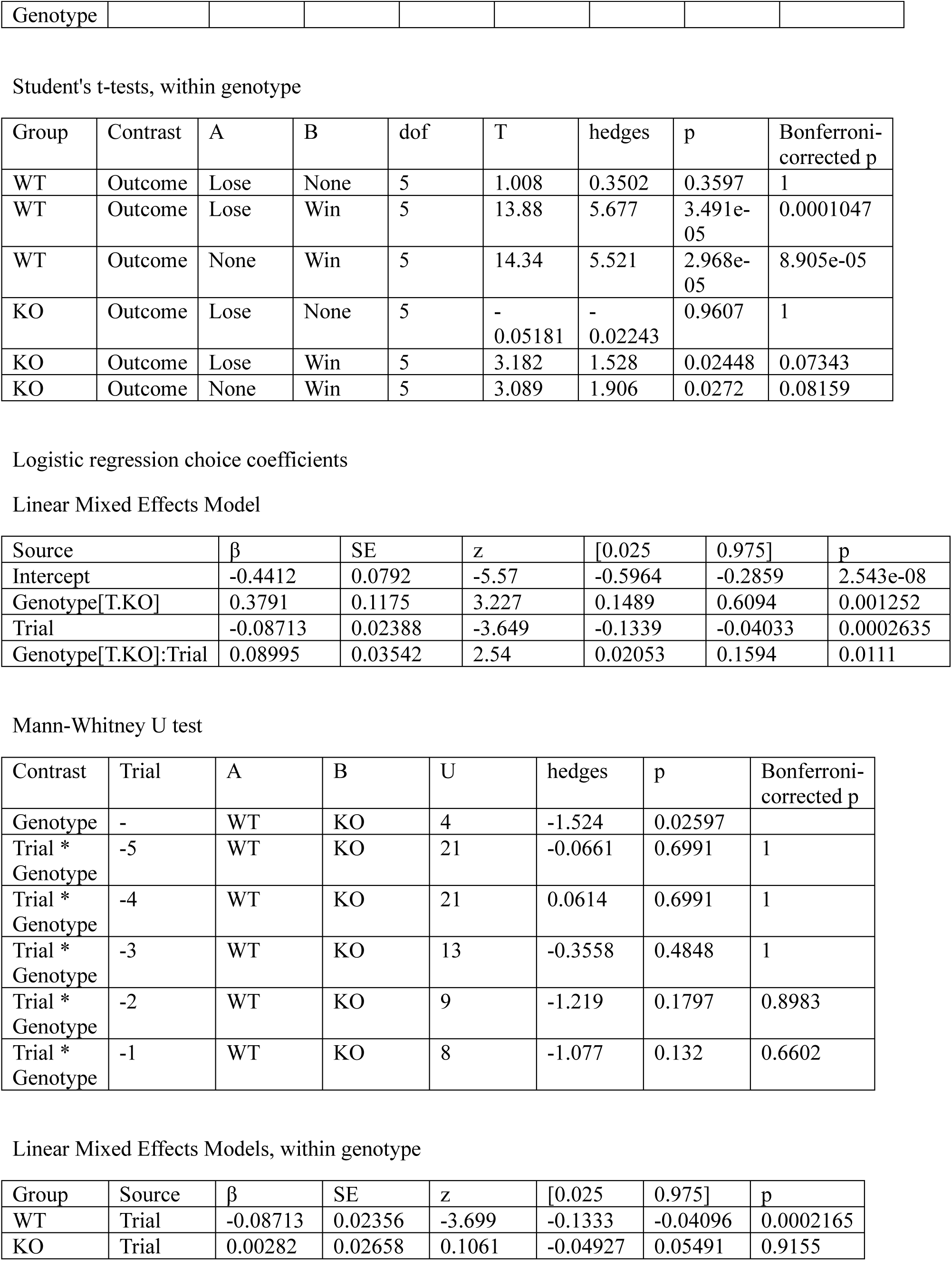
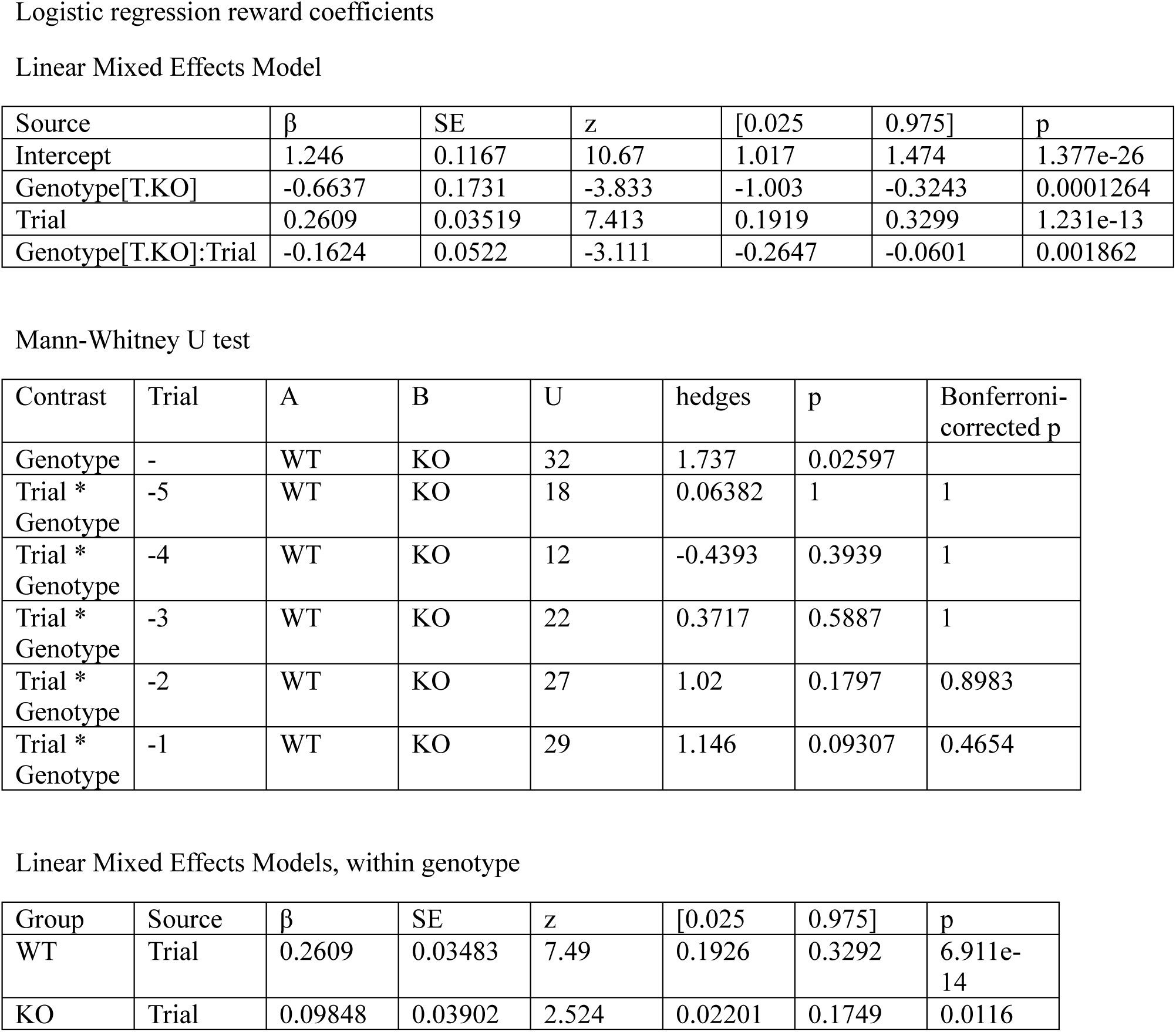

Fig 3

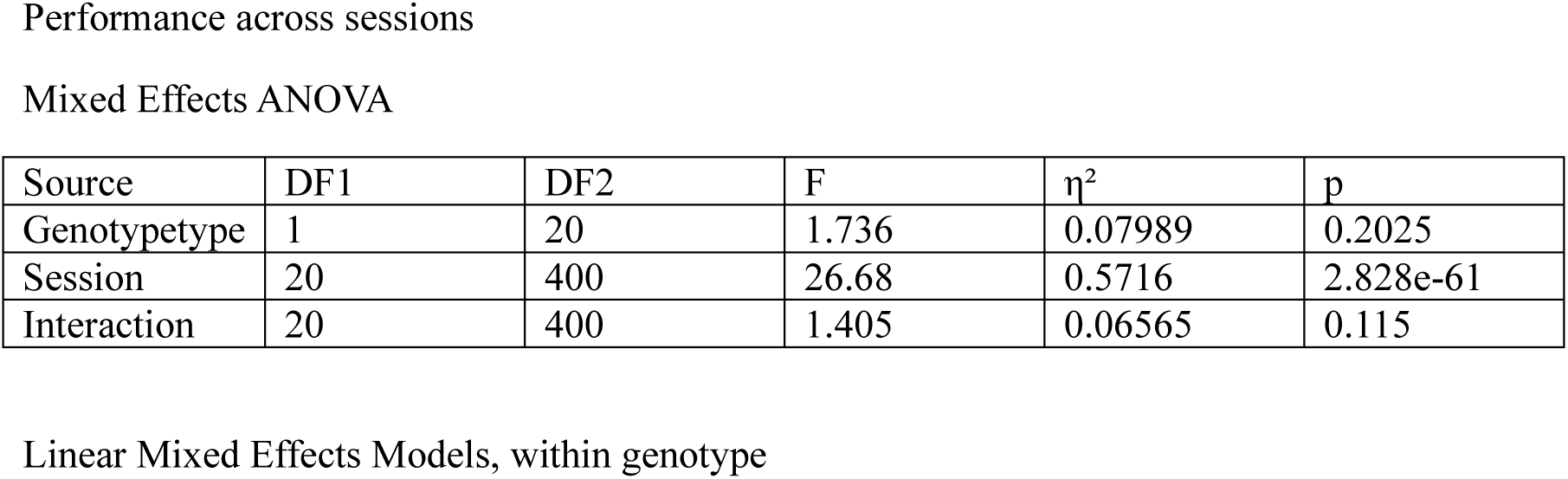
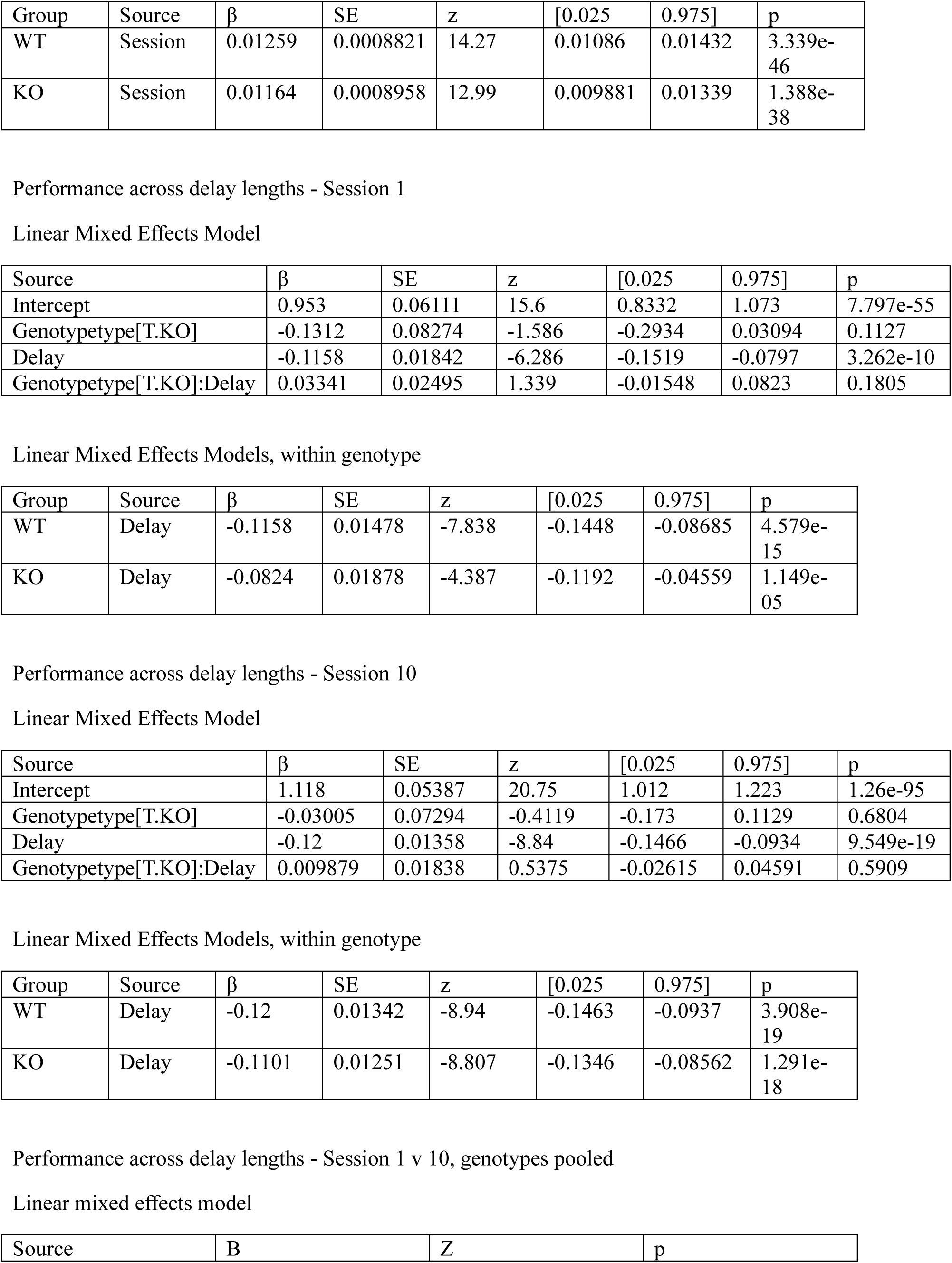
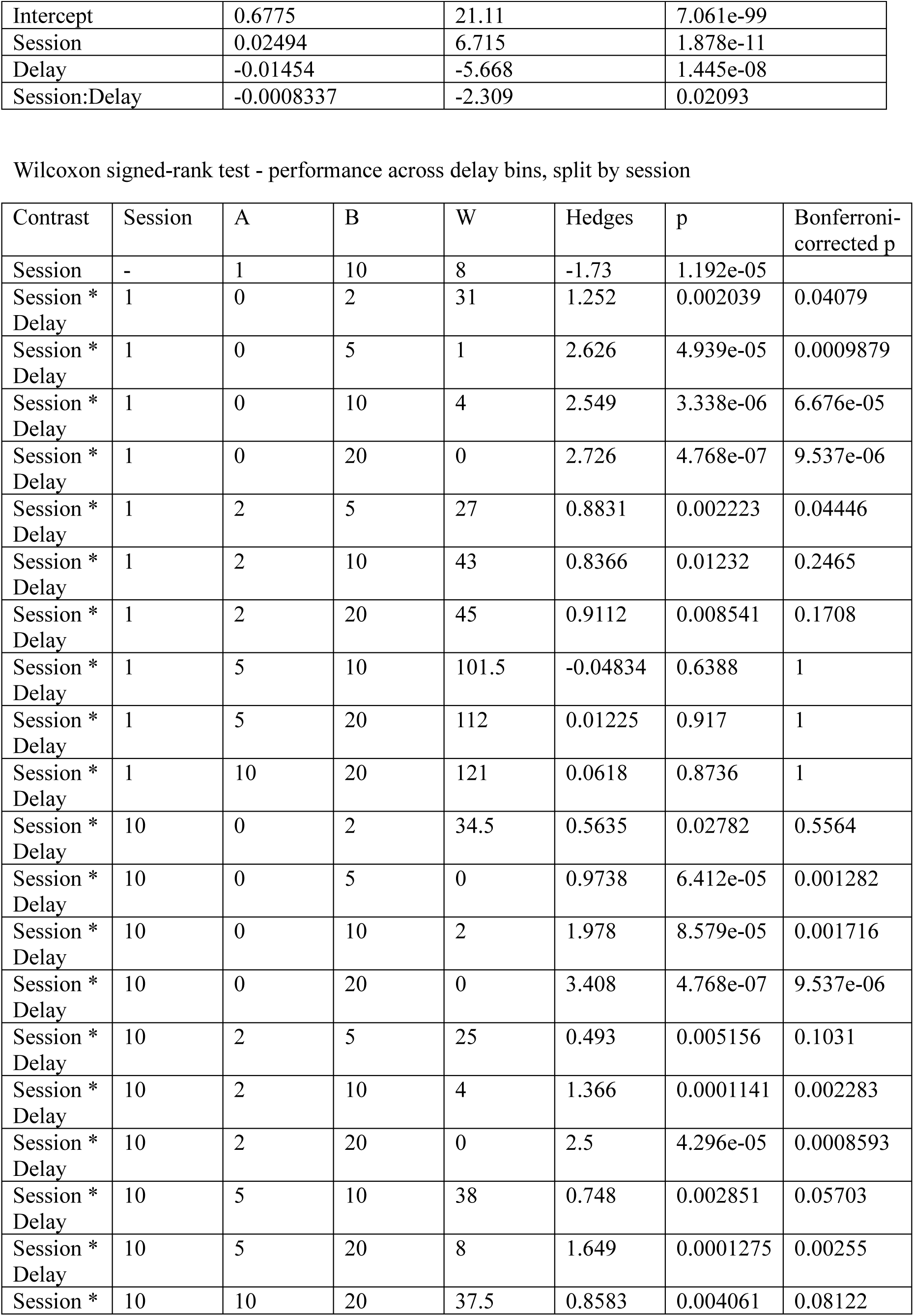
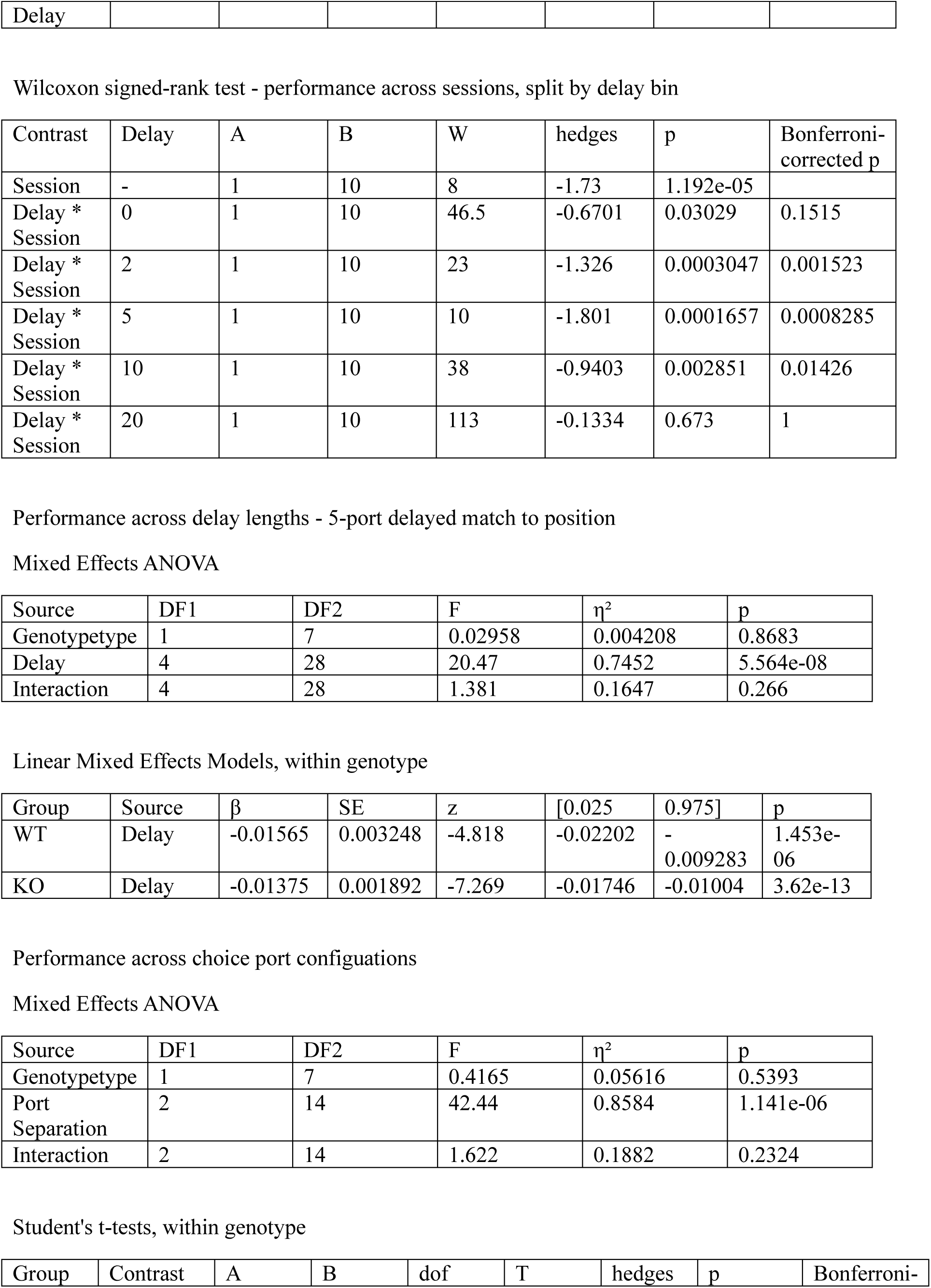
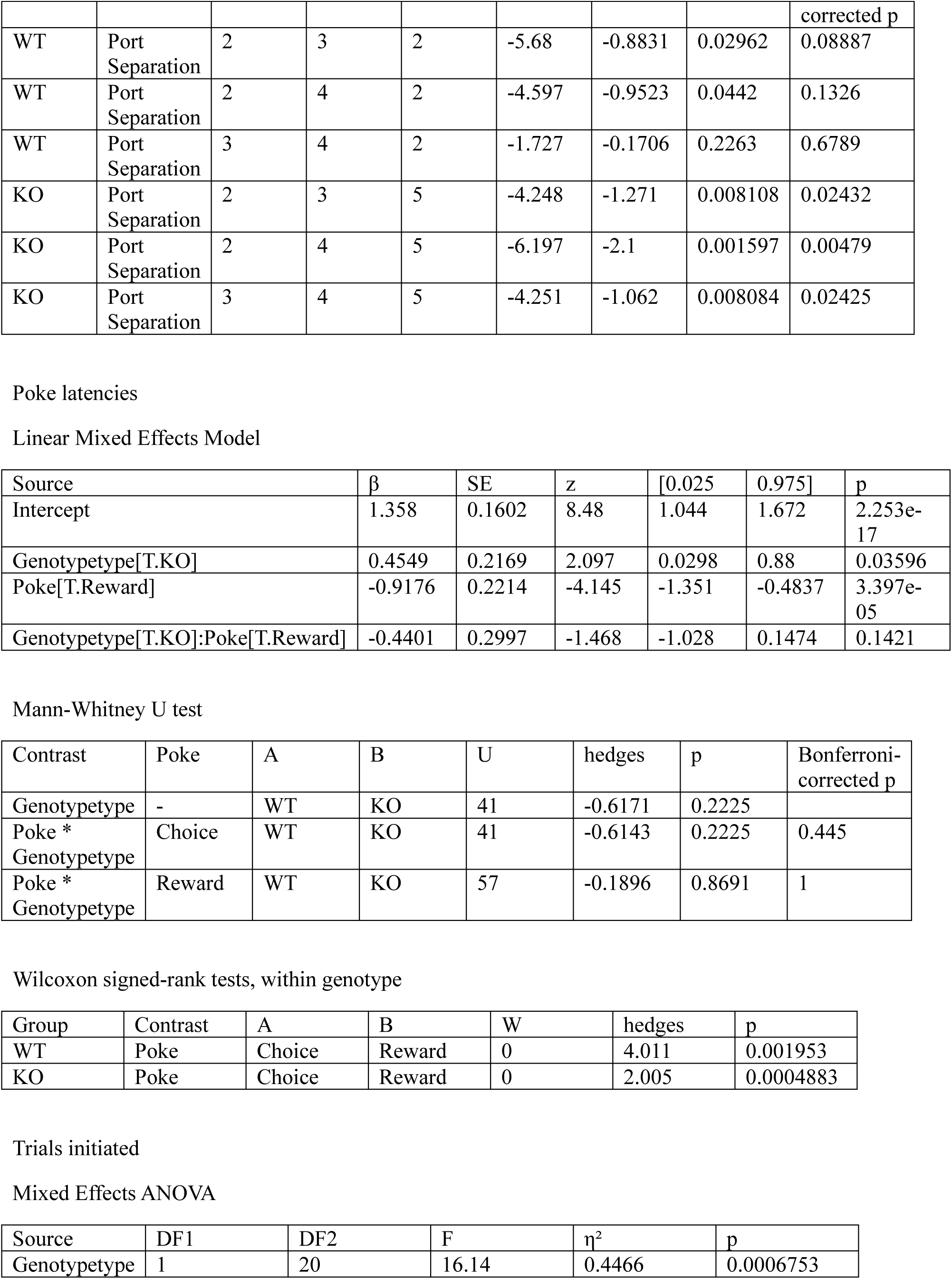
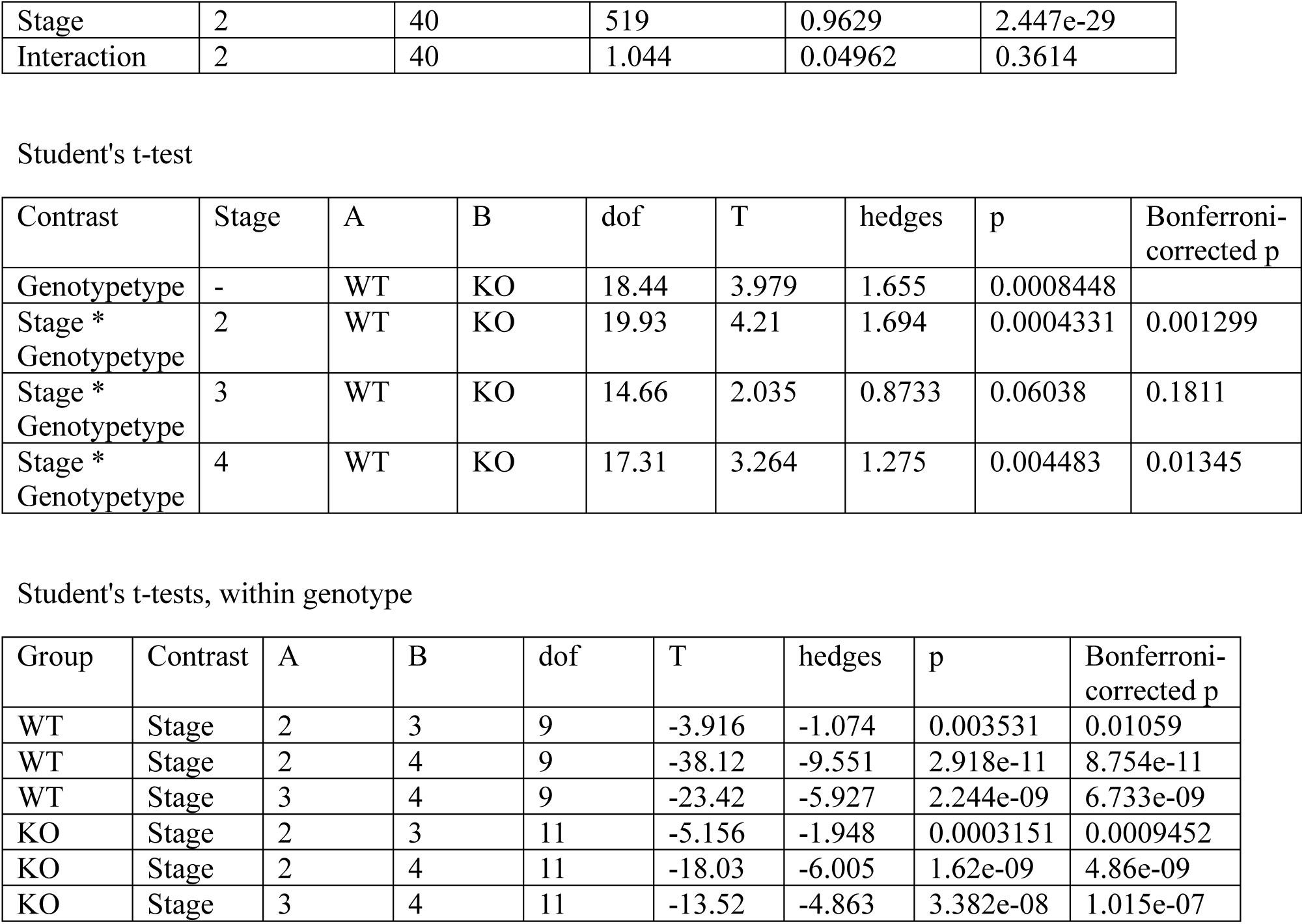

Fig 4

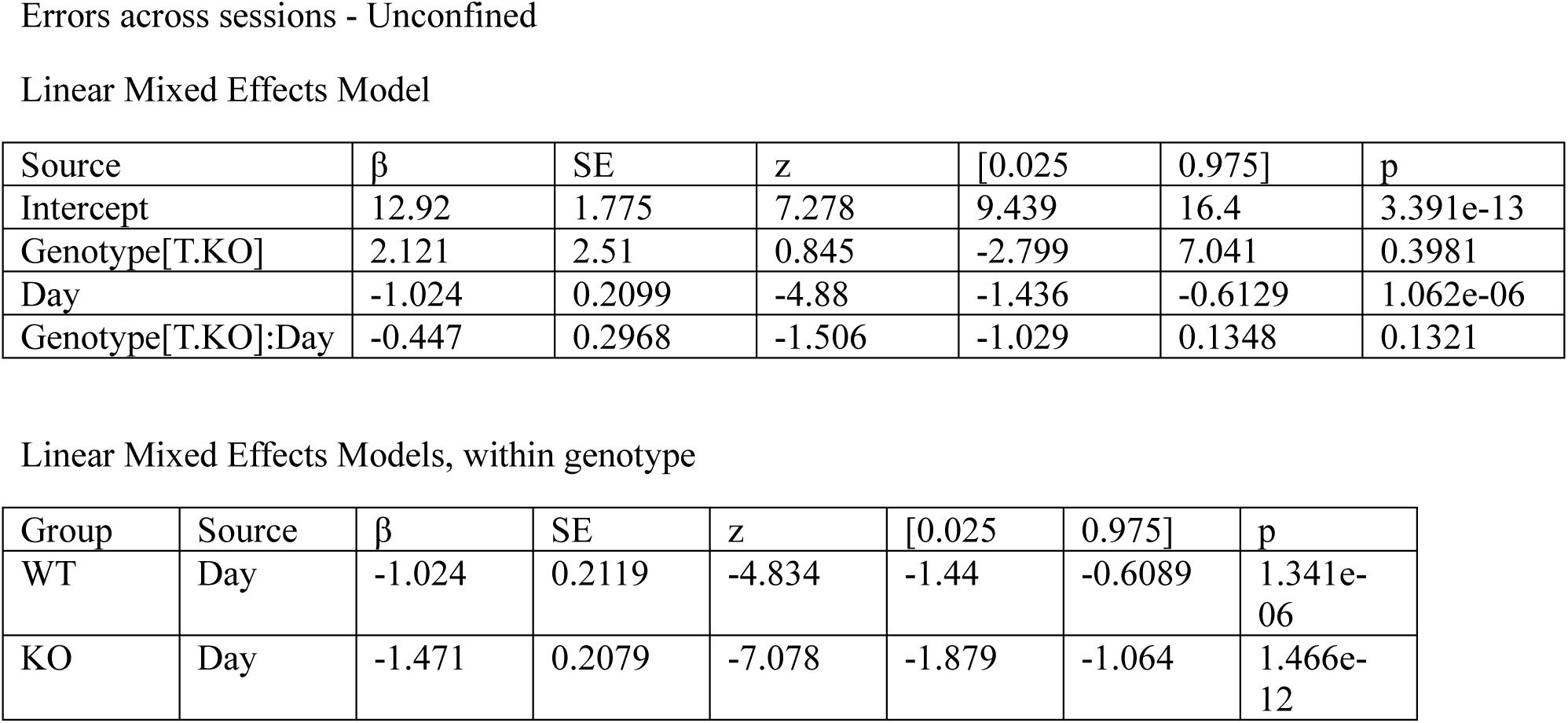
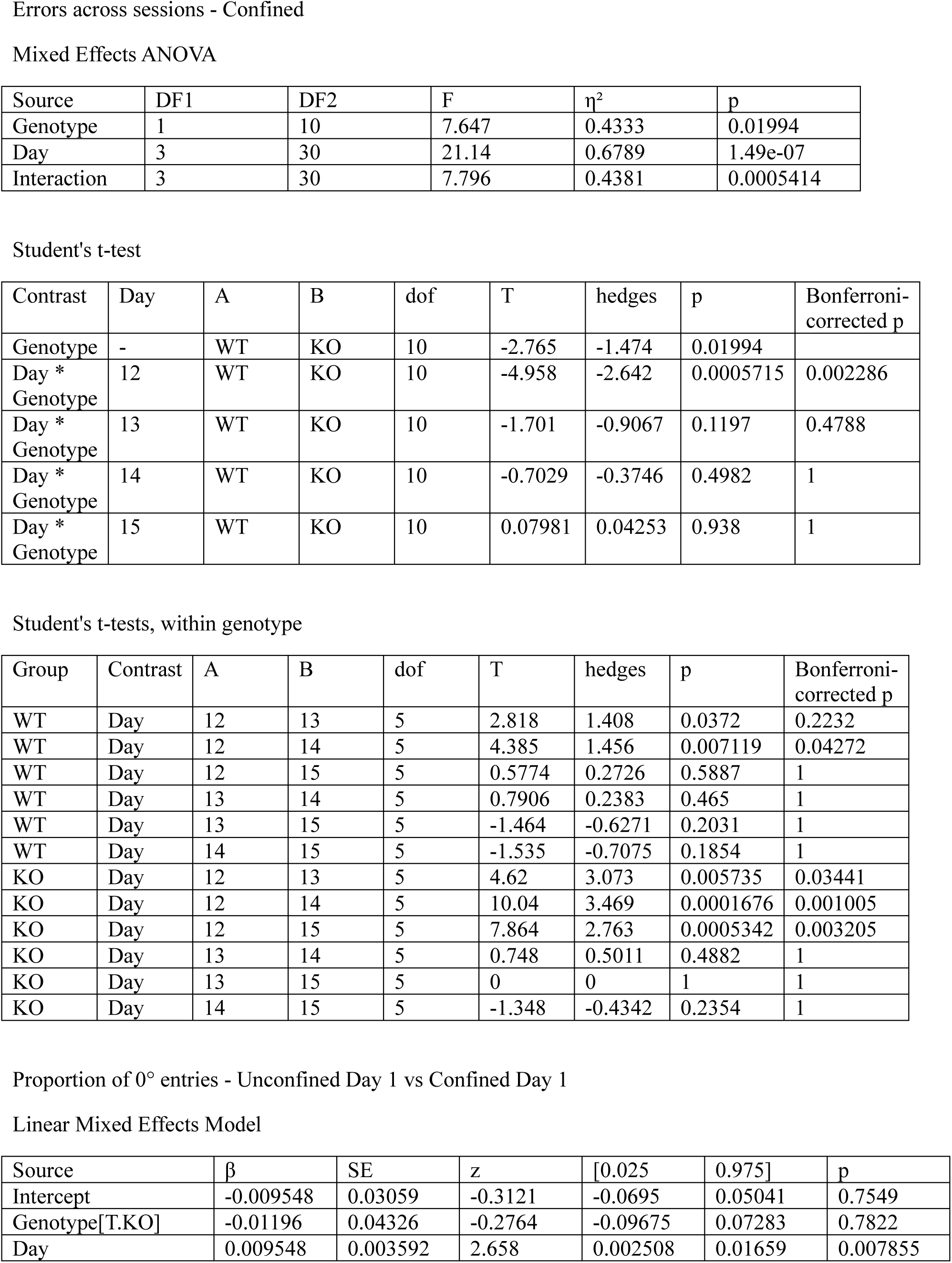
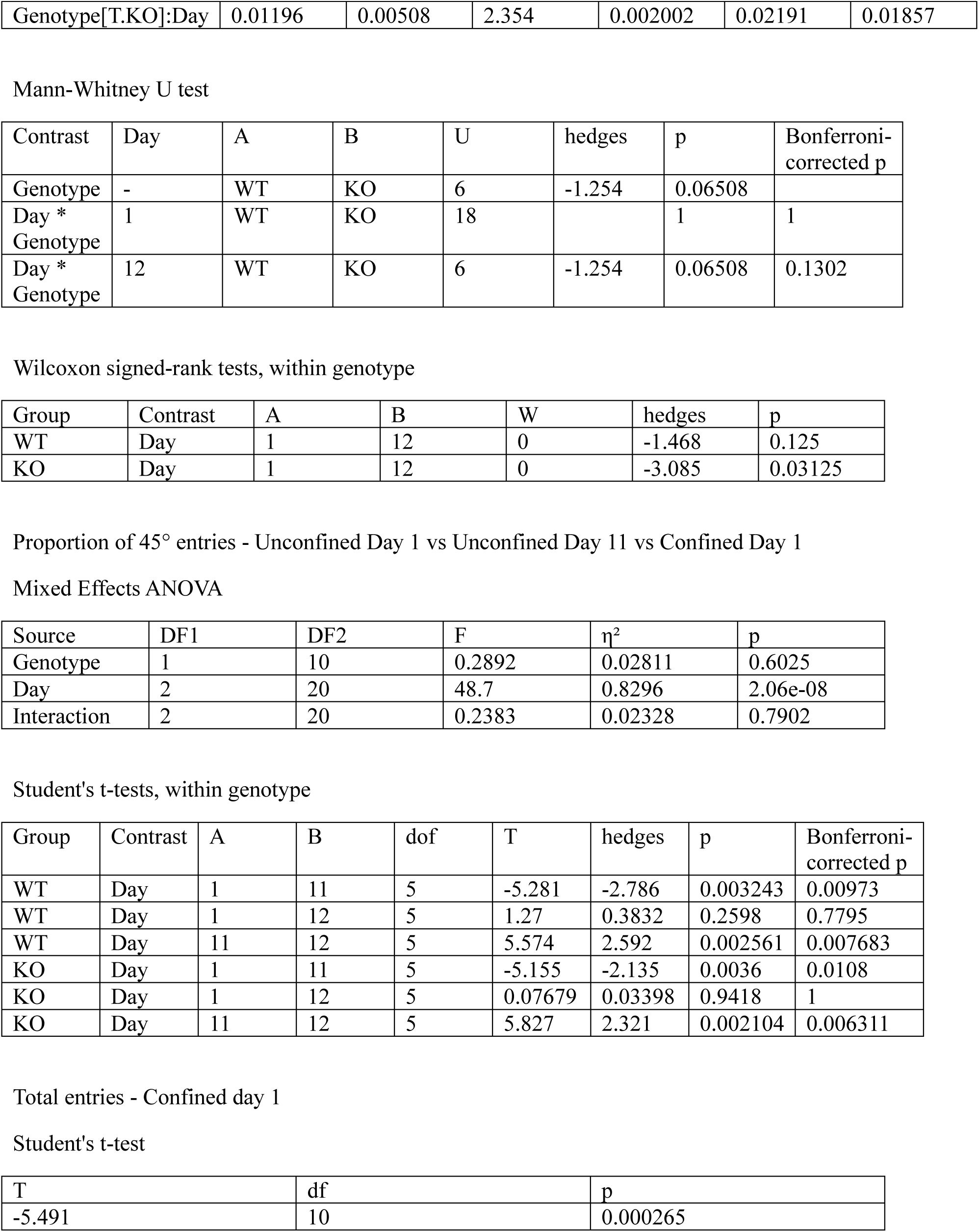
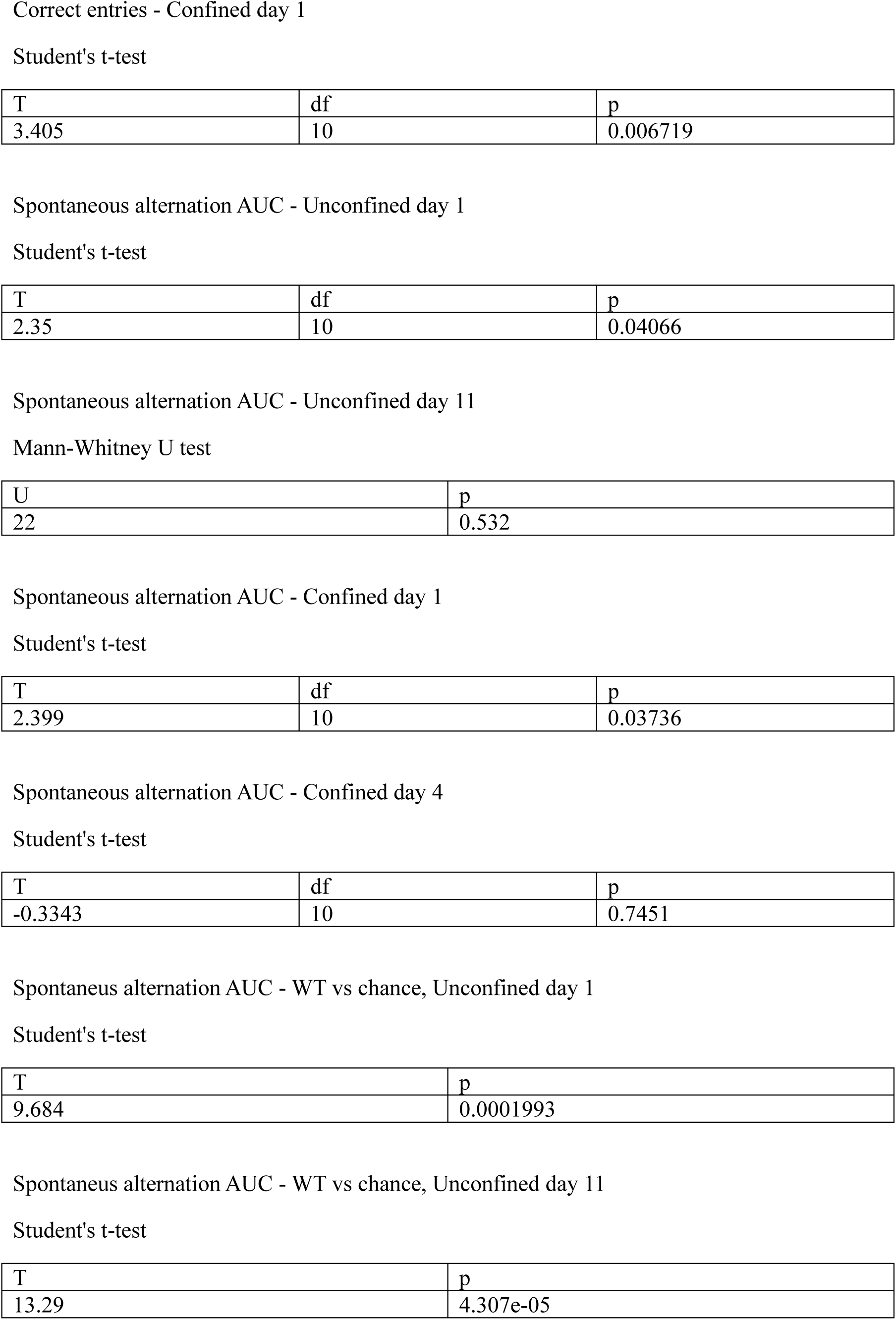
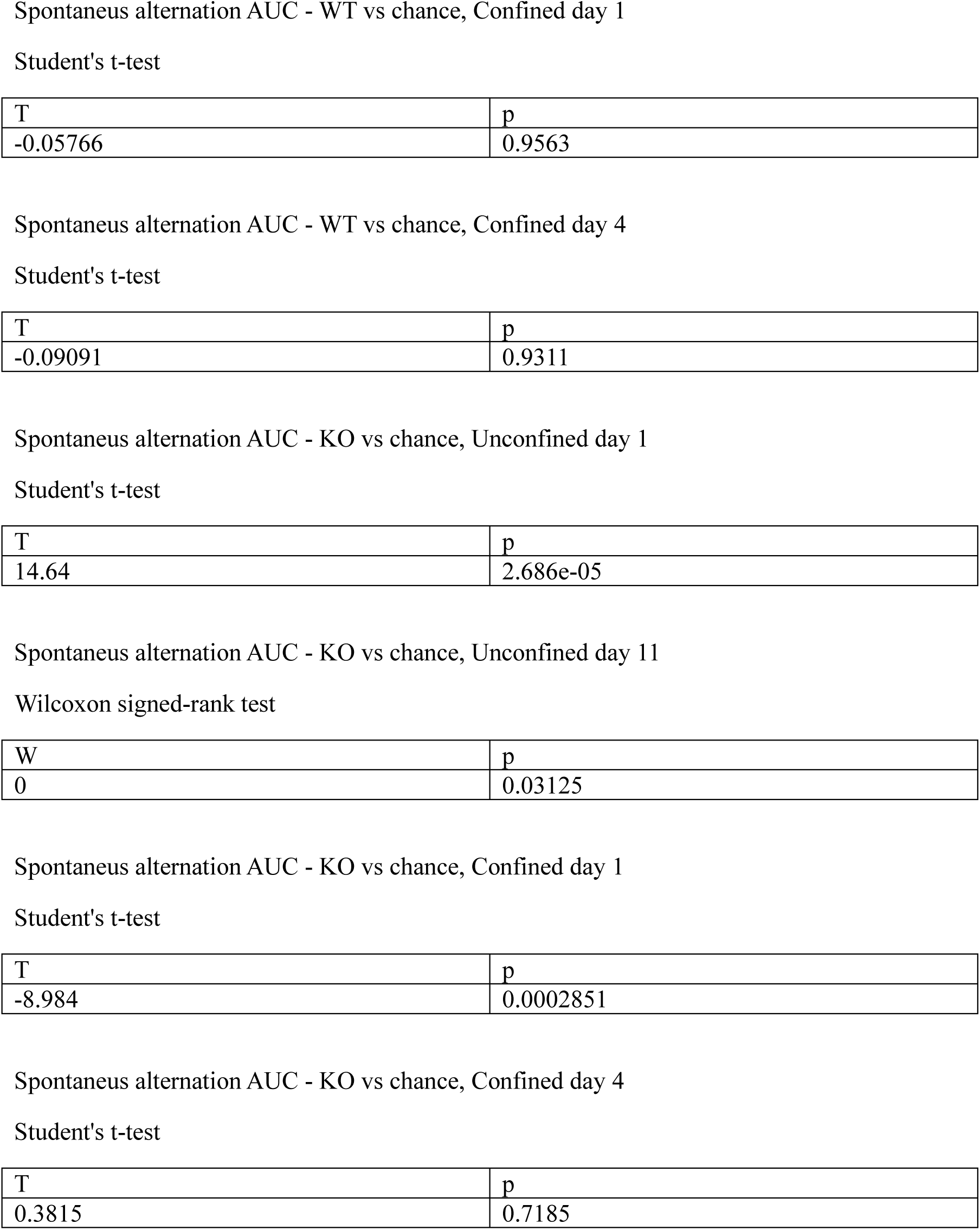

Fig 5

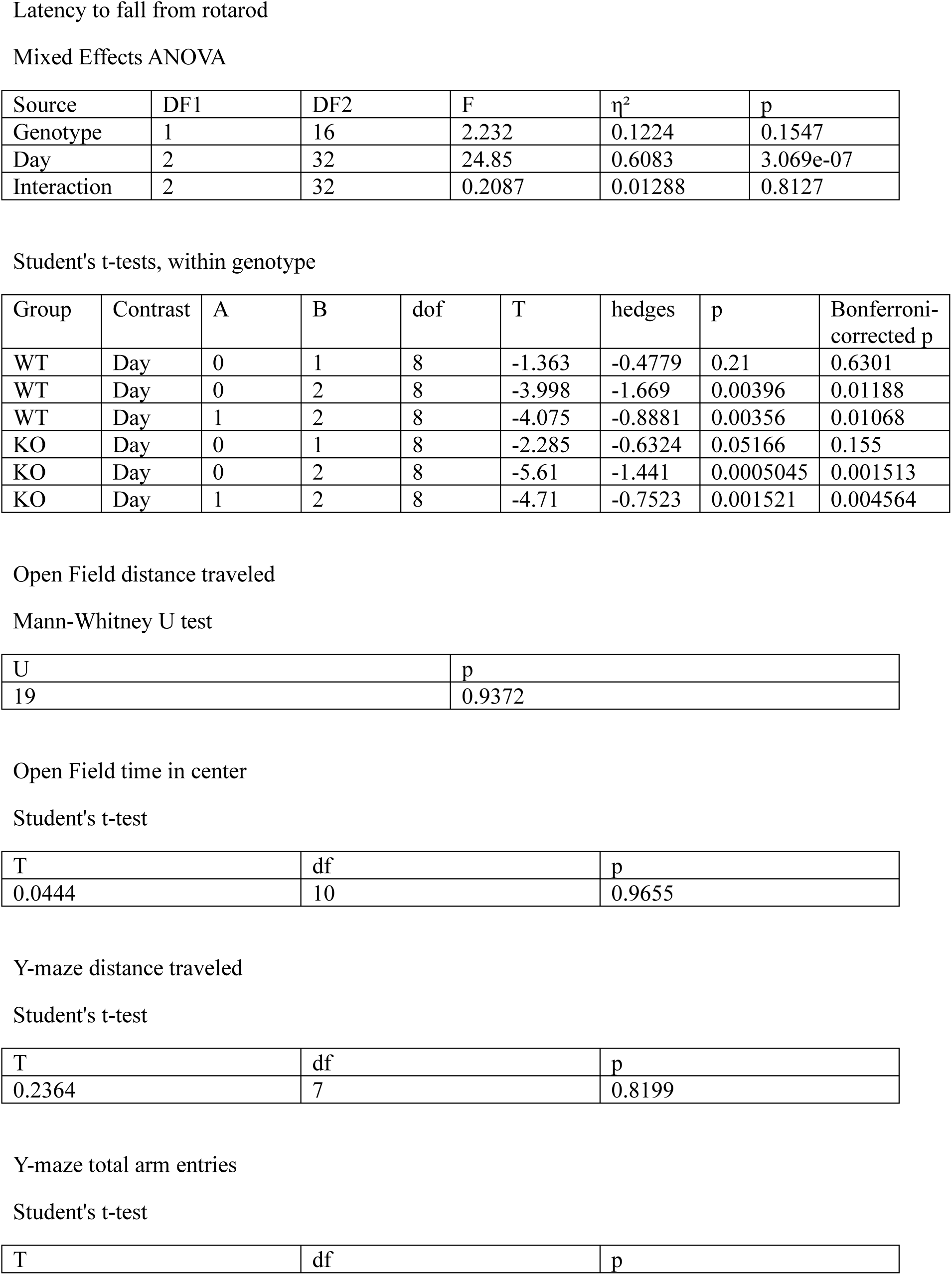
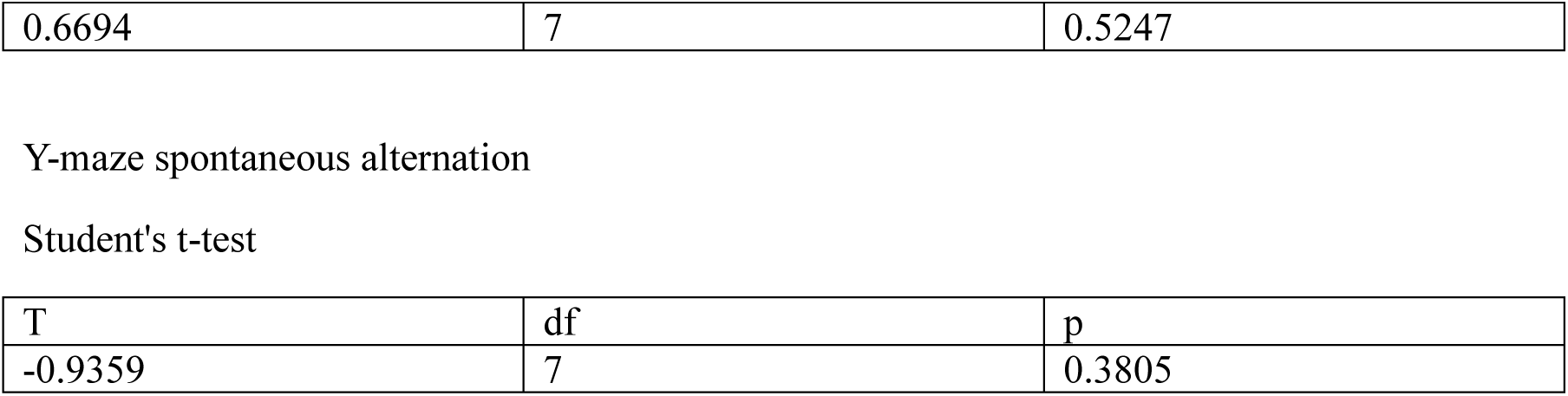

