## Supplementary Material for "Mice with impaired synaptic facilitation exhibit deficits in cognitive flexibility"

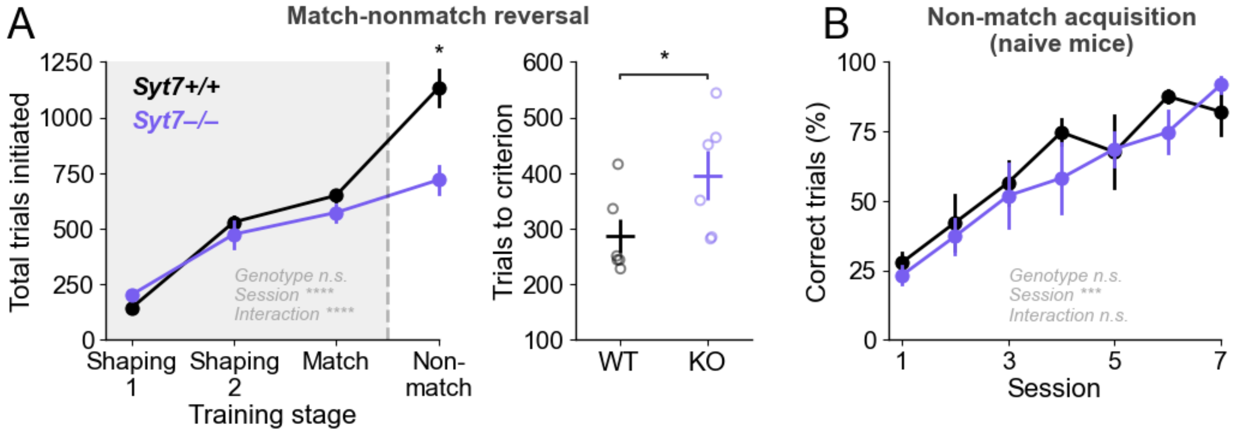


Figure 1-1: Syt7 KO show match-nonmatch reversal deficit but no impairment in nonmatch-to-sample acquisition.

A) Left: Total trials initiated across all sessions for each stage of behavioral shaping and training. Right: Total trials required to reach performance criterion (85% of trials correct).

B) Performance of mice trained on non-match to position without prior match-to-position training. Shaping sessions required mice to poke into a lit side port followed by a poke into the lit center port to receive reward. Non-match sessions required mice to sample a lit side port, poke the lit center port, choose the non-match port from the two lit side ports, and retrieve reward from the center port.

Data points are expressed as mean ± SEM, with small hollow circles representing individual animal replicates. Statistical significance is shown as *p<0.05, **p<0.01, ***p<0.001, and ****p<0.0001.

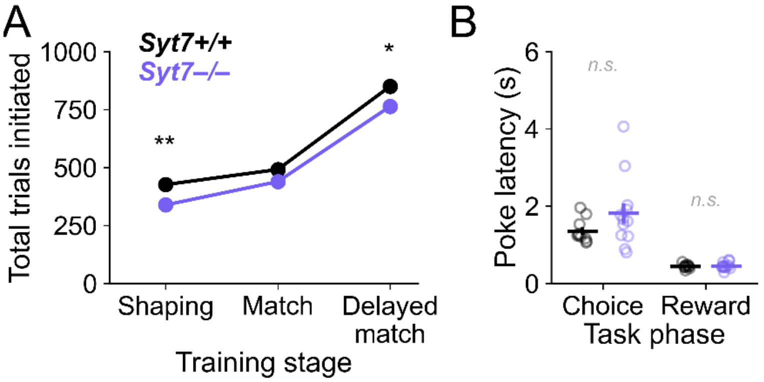


Figure 3-1: Assessment of engagement in delayed match to sample training and task.

A) Total trials initiated across training stages.

B) Average latencies to perform choice poke and reward poke during delayed match to sample task.

Data points are expressed as mean ± SEM, with small hollow circles representing individual animal replicates. Statistical significance is shown as *p<0.05, **p<0.01, ***p<0.001, and ****p<0.0001.
